# TIPPo: A User-Friendly Tool for De Novo Assembly of Organellar Genomes with HiFi Data

**DOI:** 10.1101/2024.01.29.577798

**Authors:** Wenfei Xian, Ilja Bezrukov, Zhigui Bao, Sebastian Vorbrugg, Anupam Gautam, Detlef Weigel

**Affiliations:** Department of Molecular Biology, Max Planck Institute for Biology Tübingen, 72076 Tübingen, Germany; Algorithms in Bioinformatics, Institute for Bioinformatics and Medical Informatics, University of Tübingen, 72076 Tübingen, Germany; International Max Planck Research School “From Molecules to Organisms”, Max Planck Institute for Biology Tübingen, 72076 Tübingen, Germany; Institute for Bioinformatics and Medical Informatics, University of Tübingen, 72076 Tübingen, Germany

## Abstract

Plant cells have two major organelles with their own genomes: chloroplasts and mitochondria. While chloroplast genomes tend to be structurally conserved, the mitochondrial genomes of plants, which are much larger than those of animals, are characterized by complex structural variation. We introduce TIPPo, a user-friendly, reference-free assembly tool that uses PacBio high-fidelity (HiFi) long-read data and that does not rely on genomes from related species or nuclear genome information for the assembly of organellar genomes. TIPPo employs a deep learning model for initial read classification and leverages k-mer counting for further refinement, significantly reducing the impact of nuclear insertions of organellar DNA on the assembly process. We used TIPPo to completely assemble a set of 54 complete chloroplast genomes. No other tool was able to completely assemble this set. TIPPo is comparable to PMAT in assembling mitochondrial genomes from most species, but does achieve even higher completeness for some species. We also used the assembled organelle genomes to identify instances of nuclear plastid DNA (NUPTs) and nuclear mitochondrial DNA (NUMTs) insertions. The cumulative length of NUPTs/NUMTs positively correlates with the size of the nuclear genome, suggesting that insertions occur stochastically. NUPTs/NUMTs show predominantly C:G to T:A changes, with the mutated cytosines typically found in CG and CHG contexts, suggesting that degradation of NUPT and NUMT sequences is driven by the known elevated mutation rate of methylated cytosines. siRNA loci are enriched in NUPTs and NUMTs, consistent with the RdDM pathway mediating DNA methylation in these sequences.

## Introduction

In the cells of green plants, DNA is found in three main locations: chloroplasts or chloroplast-related plastids, mitochondria, and the nucleus. The chloroplast is the primary site of photosynthesis, converting solar energy into chemical energy, while mitochondria are crucial for cellular energy metabolism. Chloroplasts and mitochondria are thought to have originated from ancient endosymbiosis events (Zimorski et al. 2014). Due to secondary and tertiary endosymbiosis, chloroplasts or plastids are present across various kingdoms, collectively referred to as photosynthetic eukaryotes (Yoon et al. 2004).

Chloroplast genomes are structurally conserved across species, and they typically comprise four distinct fragments: one large single copy (LSC), one small single copy (SSC), and two inverted repeats (IRs). In contrast, the genomes of mitochondria, present in all eukaryotic organisms except for the microorganism Monocercomonoides sp. (Karnkowska et al. 2016), vary significantly across kingdoms. The structure of bilaterian mitochondrial genomes is conserved, presenting as a single small circular DNA with sizes around 17 kb (Ladoukakis and Zouros 2017). The situation is very different in plants, which have structurally complex mitochondrial genomes with large variation in size, with the largest known mitochondrial genomes reaching up to 11 Mb (Sloan et al. 2012; Putintseva et al. 2020).

Compared to nuclear genomes, much less attention has been paid to the high-quality assembly of organellar genomes. Short read data are useful, with some caveats, for the assembly of the relatively small and conserved mitochondrial genomes of animals and chloroplast genomes of plants (Dierckxsens et al. 2017; Jin et al. 2020), but their utility is limited for the larger and more complex mitochondrial genomes of plants (Štorchová and Krüger 2024). Long and highly accurate read data have substantially enhanced our ability to assemble nuclear genomes (Wenger et al. 2019; Sereika et al. 2022). With the help of long reads, even highly repetitive regions such as centromeres and telomeres can be assembled (Naish et al. 2021; Nurk et al. 2022; Wlodzimierz et al. 2023), although challenges persist with the assembly of rDNA clusters. Moreover, the typically very high coverage of organellar genomes in data sets of genomic DNA interferes with productive assembly using standard tools, which are optimized for the nuclear genomes (Cheng et al. 2021). In addition, chloroplast and mitochondrial DNA fragments are often transferred to the nucleus, which also interferes with assembly of the true organellar genomes (Uliano-Silva et al. 2023).

Several tools have been developed to enable the specific use of long-read data for organelle genome assembly, primarily focusing on chloroplast genomes, such as Organelle_PBA (Soorni et al. 2017), ptGAUL (Zhou et al. 2023) and CLAW (Phillips et al. 2024). The general approach begins with extracting chloroplast reads from the data set by aligning long reads to the chloroplast genomes of closely related species. This is straightforward and effective for the chloroplast genome, as there are now over 12,000 published chloroplast genomes available, making it almost always possible to find a sufficiently closely related species for successful extraction of chloroplast reads. However, this approach has limitations for mitochondrial genomes, given the much smaller number of available plant mitochondrial genomes (approximately 500 as of July 2023) and the much lower conversation of mitochondrial genomes, even between closely related species. There have been ongoing efforts to assemble complex mitochondrial genomes. GSAT (He et al. 2022) begins by using short reads to construct the assembly graph and then simplifies it using long reads. However, the assembly graph created from short reads struggles to handle highly repetitive regions, making it challenging to assemble complete genomes. Recently, an alternative approach has been proposed - PMAT (Bi et al. 2024). It begins with downsampling the initial read data set to an estimated coverage of the organellar genomes that is suitable for standard assembly tools. Next, a normal assembly is performed, and then the contigs that appear to belong to organellar genomes are identified based on the presence of conserved protein coding genes. While useful, this approach may result in incomplete assemblies, especially for species with multichromosomal mitochondrial genomes where some chromosomes lack coding genes (Sanchez-Puerta et al. 2017). Clearly, the preferred approach would be a (largely) reference-free and tool for organelle genome assembly that has similar power for both chloroplast and mitochondrial genomes.

As stated above, organellar DNA can be transferred to the nucleus, and it is common to find organellar sequences in the nuclear genome (Erik Richly and Leister 2004; Hazkani-Covo et al. 2010; Michalovova et al. 2013; Zhang et al. 2020). These sequences are known as nuclear mitochondrial DNA (NUMTs) and nuclear chloroplast DNA (NUPTs). The nuclear genome evolves much faster than mitochondrial genome, typically by an order of magnitude (Wolfe et al. 1987; Drouin et al. 2008). Accordingly, NUPTs and NUMTs tend to diverge from the ancestral organellar genomes quite rapidly. By aligning NUPTs and NUMTs, which should not carry any function, to the corresponding organelle genomes, one can explore presumably neutral processes of sequence change in the integrated organellar DNA (Huang et al. 2005; Rousseau-Gueutin et al. 2011; Yoshida et al. 2014; Fields et al. 2022). Questions of interest are whether NUPTs and NUMTs behave in a similar manner, and how their evolutionary fate compares to that of other large insertions, such as transposons (Wang et al. 2013; Maumus and Quesneville 2014).

We have developed TIPPo, a user-friendly, reference-free assembly tool for plant organelle genomes that integrates TIARA, a deep-learning-based approach for organellar DNA classification (Karlicki et al. 2022), eliminating the need for knowledge of organellar genomes from closely related species genomes or nuclear genome information of the target species. We use k-mer information to optimize TIARA’s output, distinguishing NUPTs, NUMTs, and misclassifications caused by repetitive sequences. Using TIPPo, we not only successfully assembled 54 complete chloroplast genomes but also demonstrated superior performance in mitochondrial assembly compared to PMAT, revealing the complex structure of mitochondrial genomes. Additionally, we detailed the insertion patterns of NUPTs and NUMTs, and analyzed nucleotide substitutions in NUPTs and NUMTs.

## Approach

We designed and implemented a reference-free, user-friendly tool for assembly of plant organelle genomes called TIPPo from highly accurate PacBio HiFi long reads. It begins with a deep learning model to identify candidate organelle reads, followed by use of a k-mer count approach to filter out the remaining nuclear reads and finishing with assembly of the organellar genomes. **Figure 1** illustrates the entire workflow.

**Figure 1.**
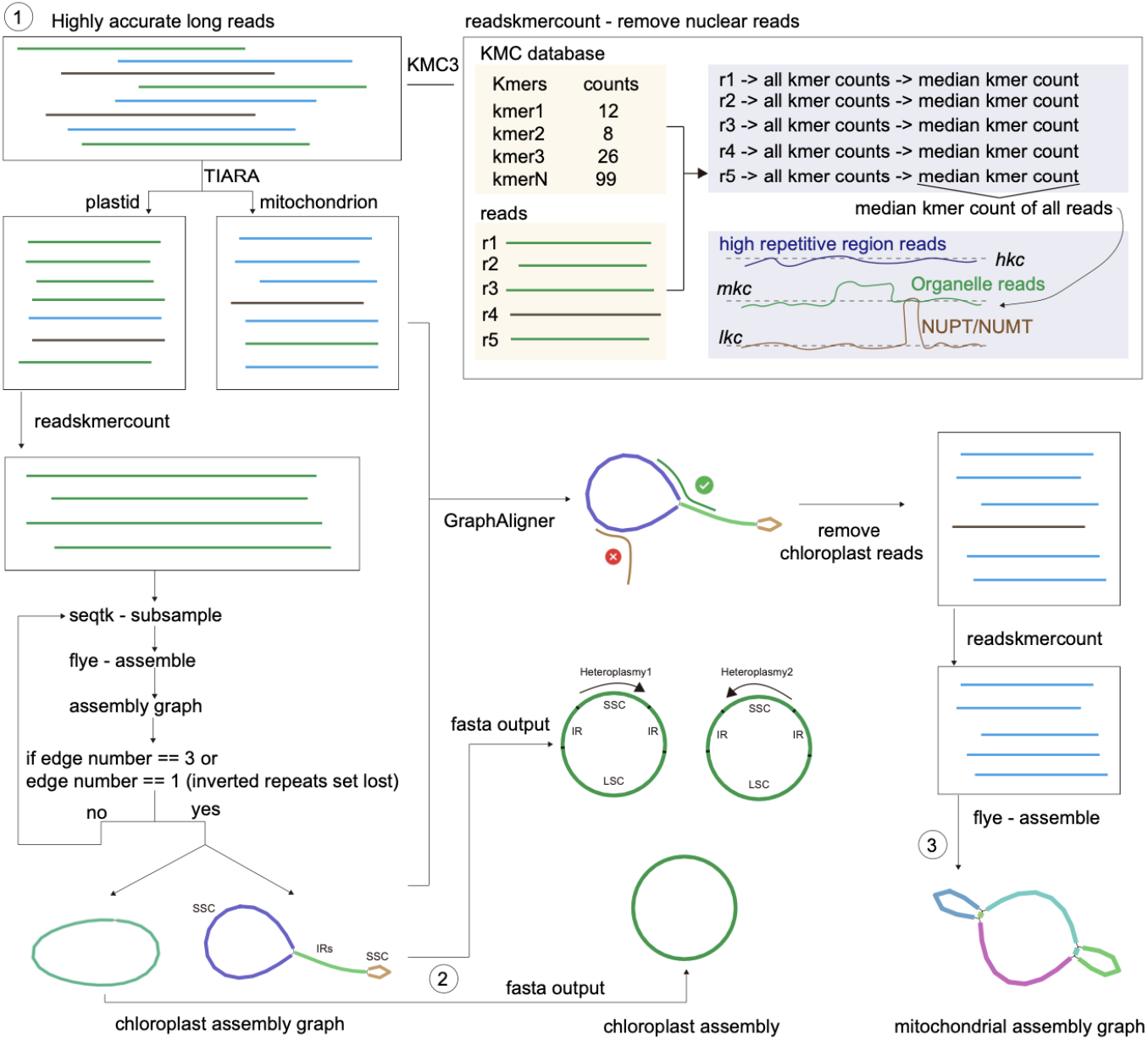
Workflow of TIPPo.

TIPPo uses TIARA (Karlicki et al. 2022)to classify the reads, a deep learning-based approach that follows a two-step process: first, it classifies reads as nuclear or organellar, and then further categorizes the organellar reads into plastid or mitochondrial. We evaluated the accuracy of TIARA (Karlicki et al. 2022) using *Arabidopsis thaliana* and *Oryza sativa* (**Figure S1, Figure S20**). As described in the original paper, TIARA will classify the NUMTs/NUPTs as organelle reads, and there is also an increased proportion of misclassification in highly repeated regions, such as centromeres and rDNA clusters. Hence, further filtering is necessary.

The assumption for subsequent filtering is that true organellar reads are the largest class in the TIARA output, and that misclassifications are relatively rare. We use KMC3 (Kokot et al. 2017) to generate a k-mer (k=31) count database from the reads identified by TIARA. Next, we perform filtering based on k-mer counts separately for chloroplast and mitochondrial reads. We use readskmercount to obtain the read median k-mer count *rmkc*, which is used as a representative for each read. Reads labeled as plastid are processed first because chloroplast genomes are more conserved than mitochondrial genomes.

After calculating *rkmc* for all input reads, the median kmer count *mkc* of all chloroplast reads, and of all mitochondrial reads after chloroplast assembly, will be used for filtering. To this end, we set the low kmer count threshold *lkc* to 0.3 * *mkc*, and the high kmer count threshold *hkc* to 5 * *mkc*. A read is removed if more than one fifth of its k-mer counts are either lower than *lkc* or higher than *hkc*. Reads with many k-mer counts below the *lkc* threshold likely originate from the nucleus, and possibly correspond to NUPTs or NUMTs. Reads with many k-mer counts above the *hkc* threshold are likely from highly repetitive nuclear regions such as centromeres, and rDNA clusters. After filtering, flye (Kolmogorov et al. 2019) is used to assemble the chloroplast genome in the first assembly step. The assembly is performed iteratively with a random selection of reads, until the assembly graph matches the typical chloroplast structure. In each assembly round, only 800 reads are used, which is around 100x coverage, since excessive coverage might negatively affect flye results. Following the assembly with flye, the assembly graph is checked for a typical chloroplast structure or a circular DNA when inverted repeats were set as lost. The structural check is aiming to match two isomeric chloroplast genomes that coexist equimolarly, differing only in the orientation of the LSC and SSC, as is the case in most land plants and algae (Palmer 1983; Aldrich et al. 1985; Wang and Lanfear 2019). Once this is achieved, the cycle ends with output of two typical heteroplasmic fasta sequences or one circular sequence.

The next step is the assembly of the mitochondrial genome. Considering that some chloroplast reads might be misclassified as mitochondrion by TIARA, GraphAligner (Rautiainen and Marschall 2020) is used to align all reads labeled as mitochondrion to the chloroplast assembly graph as a further refinement step. If the read alignment is almost end-to-end (left clip length ≤ 100 bp, right clip length ≤ 100 bp and identity > 95%), reads are considered as likely originating from the chloroplast and are removed. It is worth noting that mapping reads directly to an organelle assembly graph is the optimal solution for the organelle genome alignment, since linearized circular DNA combined with heteroplasmy will lead to clipped alignments. CLAW (Phillips et al. 2024) also addresses the alignment issues caused by a linearized circular DNA target by joining the two linear DNA sequences. Although this approach avoids clipping alignment, it introduces the issue of mapping quality of zero.

As a final step in TIPPo, the reads remaining after alignment to the chloroplast assembly graph will be processed by readskmercount to exclude reads originating from the nucleus, as described above. Given that the coverage of mitochondrion is generally lower than that of chloroplast and the genomes size are usually larger, all finally remaining reads serve directly as input to flye for generating the assembly graph.

## Results and Discussion

### Chloroplast genome assembly

Given the conserved structure of chloroplast genomes, we categorized the assemblies based on the structure on the assembly graph into three classes: 1) containing only the typical chloroplast genome or one circular DNA (complete genome); 2) consisting of the complete genome and other sequences; 3) incomplete assembly (**Figure 2**).

**Figure 2.**
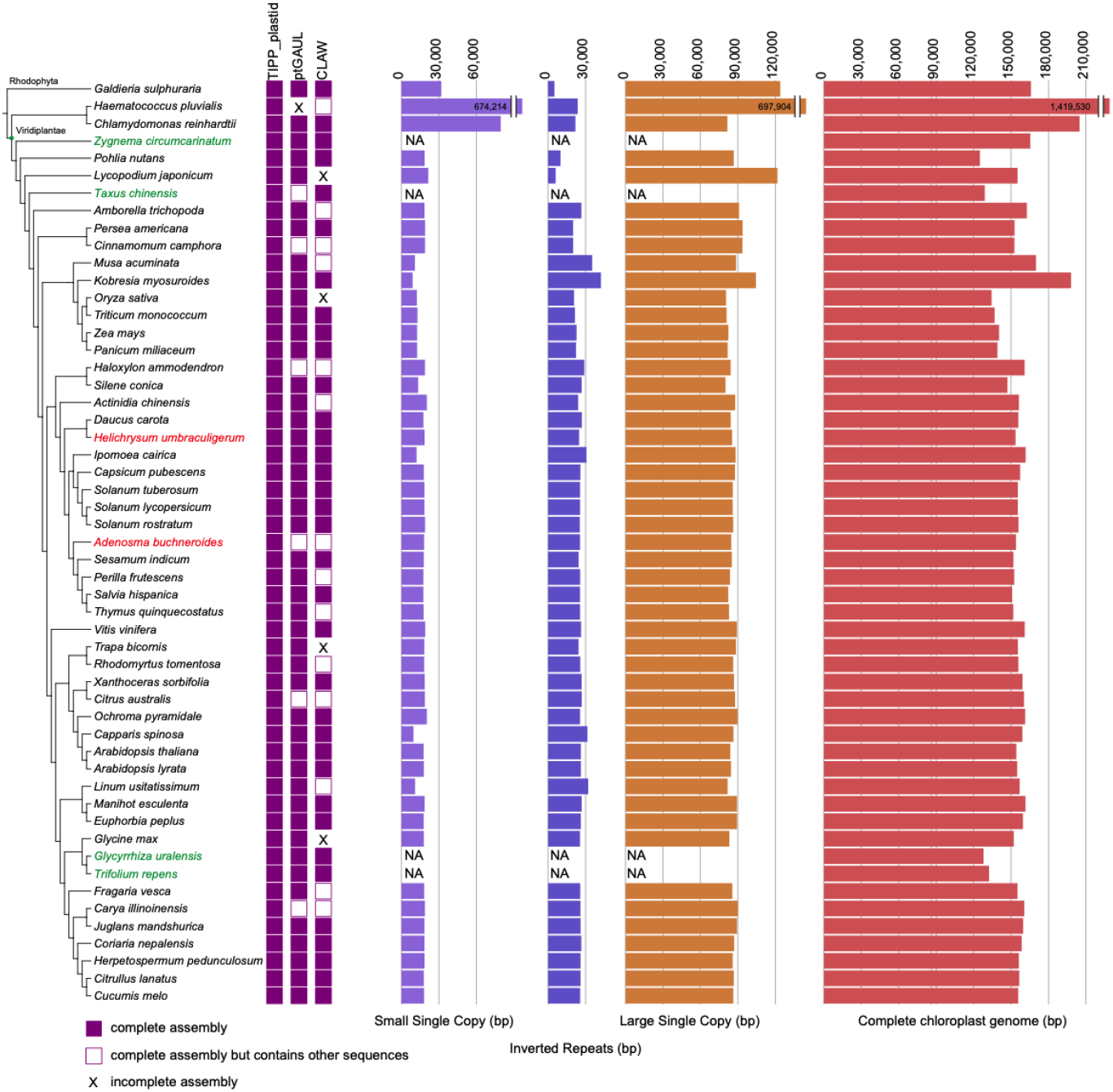
Benchmarking of four chloroplast genome assembly tools and genome statistics. See Methods for phylogenetic tree. Red species names indicate de novo assemblies for the first time. Green species names indicate loss of inverted repeats, and the three topologically defined regions are therefore not measured.

To test the performance of TIPPo, we selected 54 phylogenetically diverse planta and compared the performance with that of ptGUAUL and CLAW. Using TIPPo, we successfully assembled all 54 complete chloroplast genomes without any extraneous sequences (**Figure S2**) and assembled 48 complete chloroplast genomes at 0.5x nuclear genome coverage (**Figure S23**). We obtained two chloroplast genomes (**Figure S2**) for *Acorus gramineus*, suggesting that the sample might contain reads from two genotypes. It was therefore excluded from downstream analysis. ptGAUL assembled 46 complete genomes, produced six assemblies containing complete chloroplast genomes along with other sequences, and was unable to assemble one species (**Figure S3**). CLAW successfully assembled 35 complete chloroplast genomes, 14 assemblies included complete chloroplast genomes as well as other sequences, and it did not assemble four species (**Figure S4**).

Out of 54 species, four species have lost inverted repeats, resulting in a single circular structure in the assembly graph (**Figure S2**). For the remaining 50 species with inverted repeats, the assembly graph typically consists of three nodes: one representing the LSC, one representing the SSC, and one representing the IR, as shown in **Figure S2**. The heteroplasmy in chloroplasts is mainly mediated by an inverted repeat, so for outputs with typical chloroplast structures in TIPPo, we provide two separate configurations of the chloroplast genome.

Whole-genome alignments against published chloroplast genomes indicated high consistency between the published and TIPPo assemblies (**Figure S6**). Typical chloroplast genomes have three distinct regions: SSC, LSC and IR. Public chloroplast genomes are typically presented as linear circular DNA sequences. Thus, in whole-genome alignments, the single IR from the assembly graph aligned to two regions of the linear representations, with one forward and one reverse orientation (**Figure S6**). We also assembled two previously unpublished chloroplast genomes, *Adenosma buchneroides* (153,640 bp) (**Figure S7**) and *Helichrysum umbraculigerum* (154,011 bp) (**Figure S8**). Comparing the chloroplast genome lengths across 53 species, we observed that those from green algae are larger than those from terrestrial plants, with terrestrial plant chloroplast genomes generally around 150 kb (**Figure 2; Table S4**). The base consensus approach was also applied to *A. thaliana* and *S. conica*, resulting in only a 1 bp and 2 bp insertion, respectively, compared to the published reference genome, both occurring in homopolymer regions (**Figure S21**). These could be true minor differences between the exact germplasm used, or due to assembly errors in either the published genomes or in our assemblies.

### Mitochondrial genome assembly

Only PMAT assembled also mitochondrial genomes, and we therefore compared the ability of TIPPo to assemble mitochondrial genomes with PMAT. Given that PMAT assembled genomes often contain sequences from both organelles, we aligned distinct parts of the mitochondrial assembly graph from both PMAT and TIPPo against the chloroplast genomes assembled by TIPPo. For PMAT, mitochondrial genome assemblies from 33 out of 53 species contained also chloroplast sequences (**Figure S9**). For *Musa acuminata, Adenosma buchneroides, Trapa bicornis* (master), and *Fragaria vesca* (master), all parts aligned fully to the chloroplast genome graph, indicating that the assembly of mitochondrial genomes had failed. Since TIPPo removes chloroplast reads first, none of the assemblies contain chloroplast sequences (**Figure S9**). Thus, in subsequent analyses, we removed the chloroplast sequences from PMAT mitochondrial genome assemblies.

Given the structural diversity of plant mitochondrial genomes, it is challenging to assess the completeness of results from the assembly graph structure as we did with chloroplasts (Wang 2024). Inspired by BUSCO (Seppey et al. 2019) for assessing the completeness for nuclear genomes, we use 41 protein-coding genes collected by mitopy (Alverson et al. 2010) to evaluate the completeness of mitochondrial assemblies. Out of the 53 species, 35 mitochondrial genomes had previously been published, which we also included in our evaluation (**Table S5-S8**). Considering that the output of mitochondrial genomes from PMAT and TIPPo is in the form of assembly graphs, where large repetitive fragments are represented only once, we focused on the presence or absence of genes, and did not consider orientation or copy number.

As shown in **Figure 3A**, TIPPo and PMAT are in agreement regarding the completeness of protein-coding genes in the mitochondrial assemblies of 43 species. The results based on protein-coding genes are consistent with alignments to the published mitochondrial genomes (**Table S16**). In eight species, the mitochondrial genomes assembled by TIPPo had higher protein-coding genes completeness, while for two species PMAT outperformed TIPPo. For *Musa acuminata*, both TIPPo and PMAT failed to assemble the mitochondrial genome.

**Figure 3.**
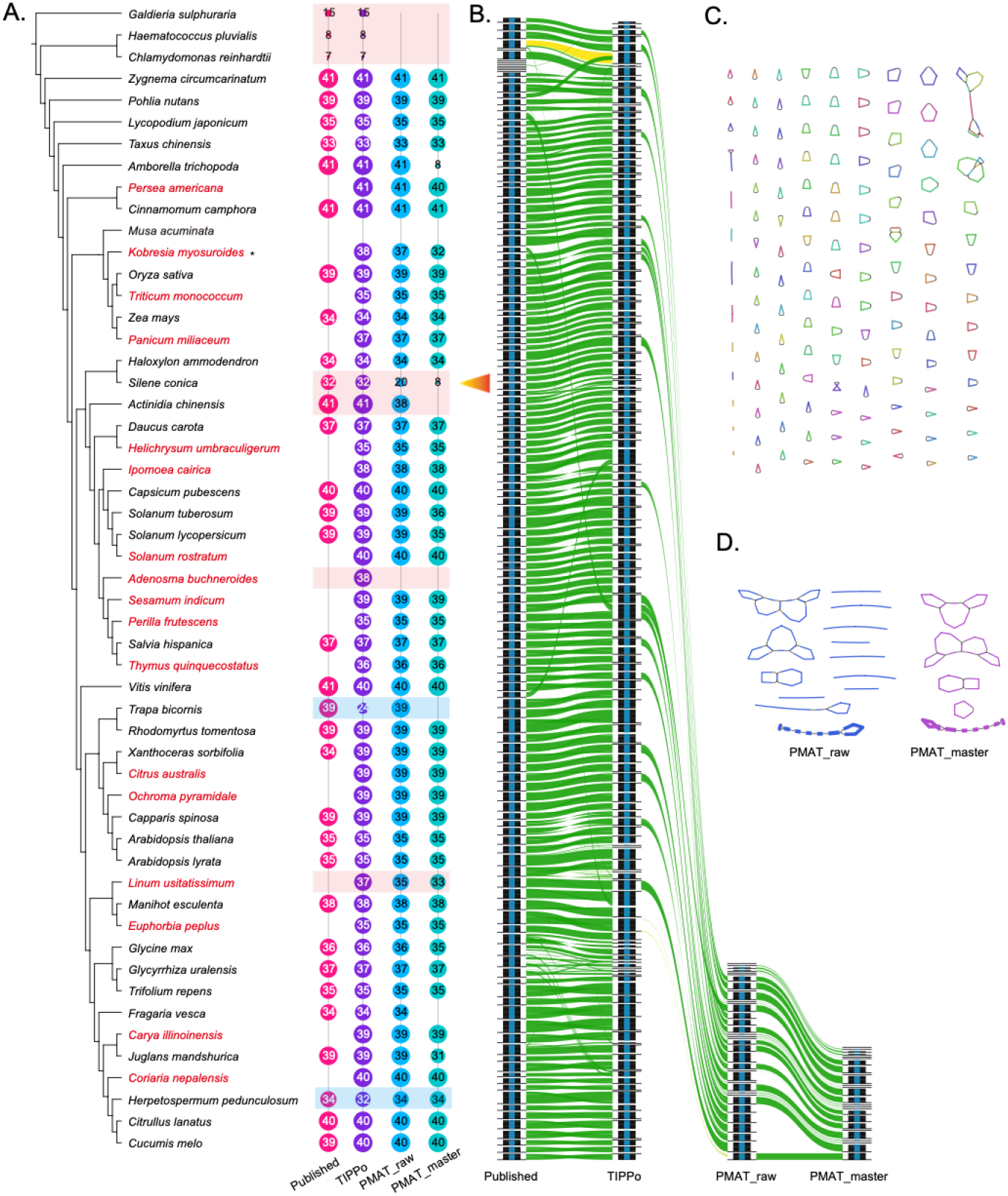
Benchmarking of mitochondrial genome assembly. A. See Methods for phylogenetic tree. Red species names indicate de novo assemblies for the first time. The numbers inside the circles indicate the number of non-redundant protein coding genes in the assembly. Light red and light blue backgrounds indicate superior results with TIPPo or PMAT. B, Whole genome alignment, including the published, TIPPo, and PMAT assemblies (both raw and master), of the *S. conica* mitochondrial genome, visualized with Alitv (v1.0.6). C. TIPPo assembly graph of *S. conica* visualized with Bandage (v0.9.0). D. PMAT assembly graph of *S. conica* visualized with Bandage (v0.9.0).

The seven species in which TIPPo was superior include one red alga and two Chlorophyta for which PMAT failed to output mitochondrial genomes, with the TIPPo assemblies matching the published assemblies for these three species. Although *Haematococcus lacustris* and *H. pluvialis* belong to the same genus, their mitochondrial genomes exhibit poor synteny (**Figure S10**).

In *S. conica*, which has one of the largest mitochondrial genomes (11 Mb), TIPPo assembled a mitochondrial genome that was highly consistent with the published genome (**Figure 3B, Figure S17**). PMAT, in contrast, only assembled parts of the mitochondrial genome. The mitochondrial assembly graph from TIPPo had numerous small circular DNAs (**Figure 3C**), which PMAT failed to identify (**Figure 3D**). A similar issue with missing small circular DNAs in PMAT occurred in *Actinidia chinensis and Linum usitatissimum*. TIPPo assembly of *A. chinensis* matched the published genome, which includes a large circular DNA of 724 kb and a smaller circular DNA of 200 kb, whereas PMAT only generates a linearized sequence of the large circle (**Figure S12**). In *L. usitatissimum*, the PMAT assembly had lost two protein-coding genes, *rpl5* and *rps14*, which are present in a circular DNA sequence assembled by TIPPo. Whole-genome alignment again indicated that PMAT the assembly had lost the circular DNA with these two genes (**Figure S13**). In *Adenosma buchneroides*, PMAT failed to assemble the mitochondrial genome, whereas TIPPo assembled a 346 kb linear DNA sequence containing 38 protein-coding genes (**Figure S14**). Given the number of protein-coding genes in related species — 39 in *Sesamum*, 35 in *Perilla*, 37 in *Salvia*, and 36 in *Thymus* — this suggests that the linear DNA sequence from TIPPo is largely complete. When using low sequence depth data as inputs for the assemblies (1x and 0.5x nuclear genome coverage), both PMAT and TIPPo showed varying degrees of incomplete assembly. Therefore, for assembling mitochondrial genomes, we do not recommend using ultra-low coverage (**Figure S22**).

As mentioned, PMAT outperformed TIPPo for two species. For *Trapa bicornis*, the TIPPo assembly graph comprised only linear DNA fragments, indicating the erroneous identification of a large number of non-mitochondrial reads. Using verkko to construct a whole-genome assembly graph revealed that *Trapa bicornis* possesses a large rDNA cluster that is misidentified by TIARA (**Figure S15**). For *Herpetospermum pedunculosum*, the TIPPo assembly lacked two genes, *nad3* and *atp6*, due to over-filtering by the k-mer approach (**Figure S19**). However, the PMAT raw assembly included non-mitochondrial fragments (**Figure S16**).

### Computational cost

Using data from 53 species, we performed chloroplast genome assembly with three different tools: TIPPo (chloroplast mode), ptGAUL, and CLAW, all with default parameters. Our results show that both TIPPo and CLAW are approximately five times slower than ptGAUL (**Figure 4A**). Regarding peak memory usage, TIPPo required the most memory, consuming three times more than CLAW and five times more than ptGAUL (**Figure 4B**). For mitochondrial genome assembly, we utilized PMAT in mt mode and TIPPo in organelle mode. PMAT was approximately eight times slower than TIPPo and consumed four times more memory (**Figure 4A and 4B**). For detailed time and memory usage at the different coverages used, please refer to **Table S9-10**.

**Figure 4.**
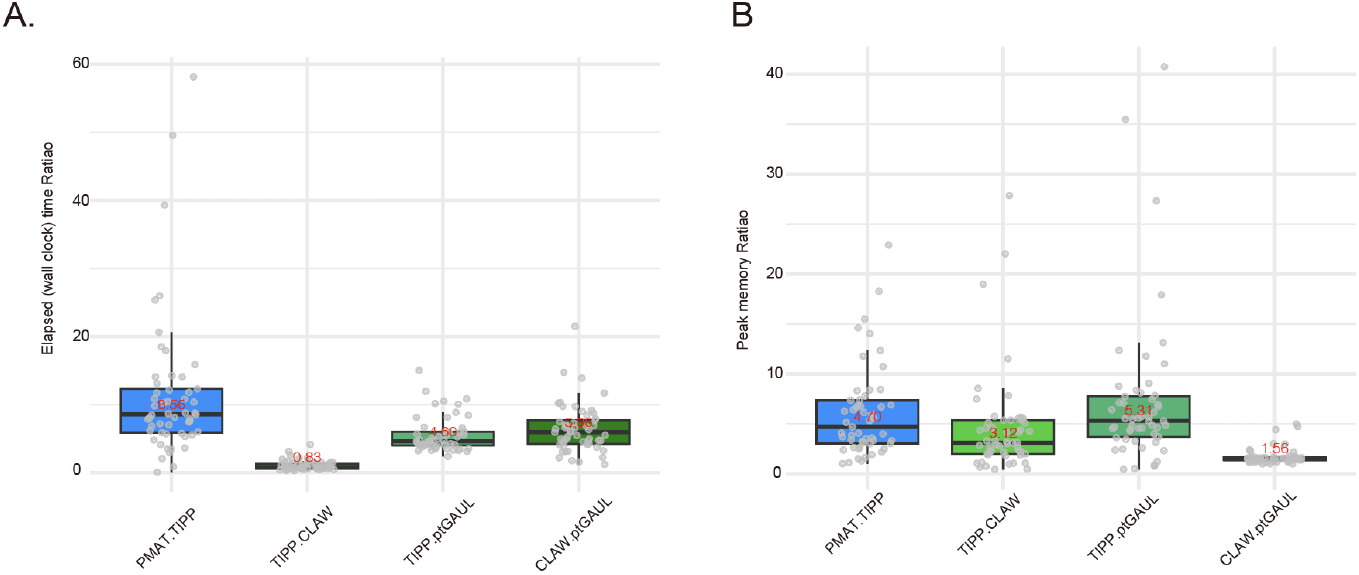
Computational cost. A. Ratio of elapsed times between each pair of the four tools. B. Ratio of peak memory usage between each pair of the four tools. Grey dots indicate different species. The means are shown as horizontal lines, with the upper and lower box indicating the interquartile range (IQR), and the whiskers extending to the most extreme values within 1.5 times the IQR from the first and third quartiles.

### Identification of NUPTs/NUMTs

Next, we wanted to know whether we could improve on the accurate identification of NUPTs and NUMTs and the elimination of potential contamination of nuclear assemblies with pieces of organellar genomes. High-quality nuclear genomes assembled from PacBio HiFi data are available for all of the species used in this study except *Lycopodium japonicum, Ochroma pyramidale* and *Perilla frutescens*, with the assemblies of the latter two being highly fragmented. Because algal genomes are small and have very few NUPTs and NUMTs (Zhang et al. 2020), we excluded them from further analysis. *Musa acuminata* was not included either, because we had not been able to assemble the mitochondrial genome. For all other 45 nuclear genome assemblies, we retrieved all contigs/scaffolds over 500 kb.

The species with the longest cumulative lengths of NUMTs were *S. conica, Amborella trichopoda, Triticum monococcum, Capsicum pubescens* and *Taxus chinensis*. This might be attributed to *S. conica* and *A. trichopoda* having large mitochondrial genomes (11 and 3.9 Mb) and *T. monococcum, C. pubescens*, and *T. chinensis* having large nuclear genomes (5, 3.9 and 10 Gb). The latter three species also had the highest cumulative lengths of NUPTs (**Table S11**). As observed before (Zhang et al. 2020), both NUPT and NUMT lengths are positively correlated with nuclear genome size in plants (Pearson’s correlation coefficients of 0.63 and 0.56) (**Figure 5 A&B**). However, no significant association was found when comparing species across different kingdoms (E. Richly and Leister 2004). Since NUPTs and NUMTs are part of the nuclear genome, their lengths are also positively correlated (**Figure 5C**).

**Figure 5.**
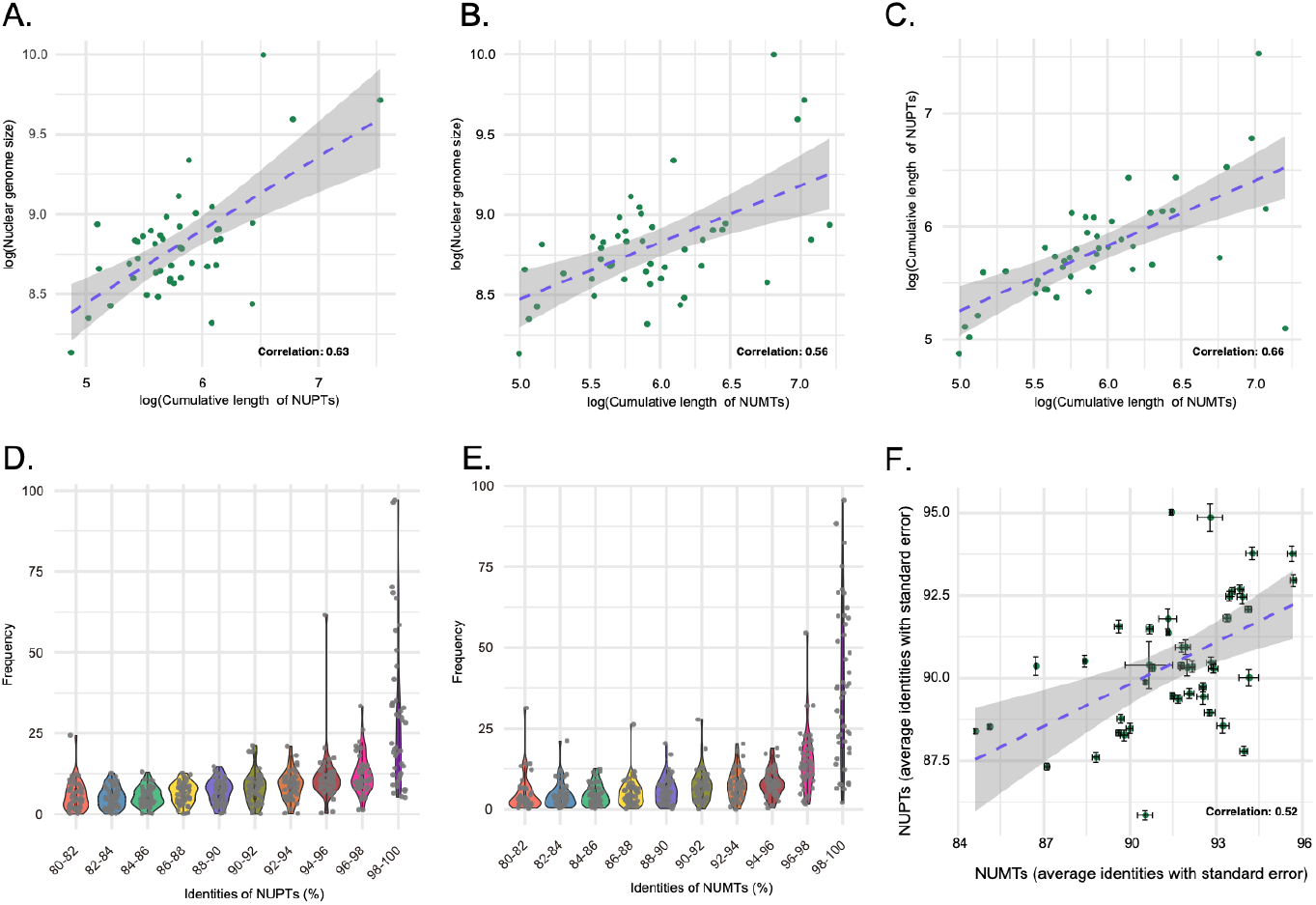
Comparison of NUPT and NUMT sequences and the corresponding organellar genomes. A. Comparison of cumulative lengths of NUPTs and of nuclear genome size. B. Comparison of cumulative lengths of NUMTs and of nuclear genome size. C. Comparison of cumulative lengths of NUPTs and of NUMTs. D. Cumulative length distribution of NUPTs across different identities. E. Cumulative length distribution of NUMTs as a function of sequence identity with the corresponding mitochondrial genome. F. Correlation between NUPT/chloroplast genome identity and NUMT/mitochondrial genome identity. Bars indicate standard errors.

NUPTs and NUMTs appear to evolve mostly neutrally, as evidenced by the gradual accumulation of mutations (Huang et al. 2005; Noutsos et al. 2005). Because the substitution rates of plant organellar genomes is typically an order of magnitude lower than that of nuclear genomes (Wolfe et al. 1987; Drouin et al. 2008), the number of differences between NUPT and NUMT sequences and the corresponding organellar genomes reflect the age of nuclear insertions (Erik Richly and Leister 2004; Michalovova et al. 2013; Yoshida et al. 2019). We found that recent insertion events, with sequence identities of 98% to 100%, are most frequent (**Figure 5D, 5E, Table S12**), which is also reflected by the correlation of average sequence identities between NUPTs and NUMTs and their organellar genomes being well correlated (Pearson’s correlation coefficient = 0.52) (**Figure 5F**). We conclude that NUPTs and NUMTs tend to degrade rapidly, which is consistent with individual NUPTs and NUMTs in *A. thaliana* genomes having low allele frequencies (Igolkina et al. 2024).

### Substitution spectra of NUPTs/NUMTs

C:G>T:A substitutions dominate the substitution spectrum in *A. thaliana* mutation accumulation lines, both in the greenhouse and the wild, although not in older natural populations (Ossowski et al. 2010; Cao et al. 2011; Exposito-Alonso et al. 2018; Weng et al. 2019). The excess of C:G>T:A substitutions has been attributed to spontaneous deamination of methylated cytosines (Ossowski et al. 2010), which is found in plants in three contexts, CG, CHG and CHH, with most of it in the CG context (Law and Jacobsen 2010). Previous studies have found that C:G>T:A substitutions to be the most common substitutions in NUPTs and NUMTs (Huang et al. 2005; Rousseau-Gueutin et al. 2011; Fields et al. 2022). We confirm this phenomenon in our set of 45 species, with the highest substitution rates at CG sites (**Figure 6, Tables S13, S14**).

**Figure 6.**
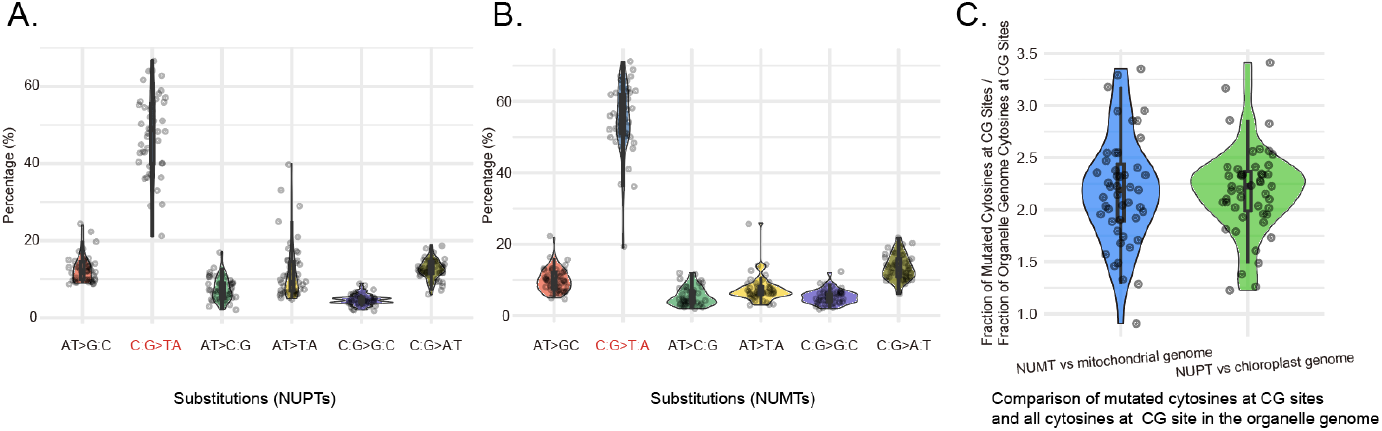
The landscape of substitutions in NUPTs and NUMTs. A. Distribution of nucleotide substitutions in NUPTs, inferred from sequence comparison with the corresponding chloroplast genome. B. Distribution of nucleotide substitutions in NUMTs, inferred from sequence comparison with the corresponding mitochondrial genome. C. Enrichment of cytosine substitutions in NUPTs and NUMTs at CG sites.

### siRNA targeting NUPTs and NUMTs

The increased substitution rate at CG sites in NUPTs and NUMTs suggested that these are often methylated, which has been directly confirmed in several instances (Yoshida et al. 2014; Fields et al. 2022). The most common type of DNA methylation in plants, RNA-directed DNA methylation (RdDM), is associated with small interfering RNAs (siRNAs) (Sigman and Slotkin 2016), and we therefore tested the hypothesis that NUPTs and NUMTs are enriched for siRNAs. In a previous study, siRNA data were generated for 11 of the 45 species that we investigated (Lunardon et al. 2020), and we annotated siRNA loci by mapping siRNA reads (Axtell 2013).

For all 11 species, the overlap of siRNA loci with NUPT/NUMTs was significantly higher than expected by chance (**Figure 7, Table S15**), demonstrating that siRNAs are indeed enriched in NUPTs and NUMTs.

**Figure 7.**
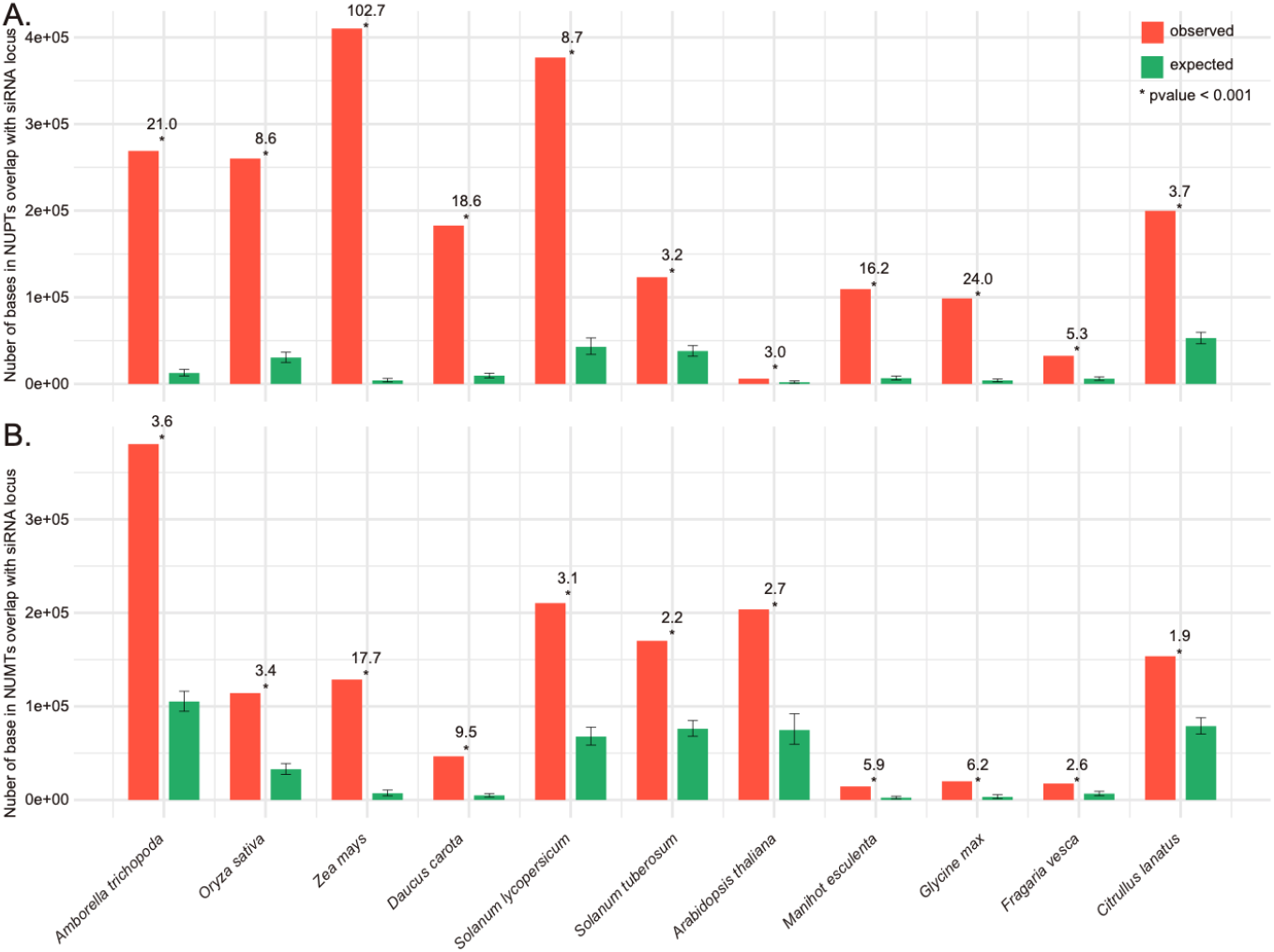
Enrichment of siRNAs in NUPTs and NUMTs. A. Overlap of siRNA loci with NUPTs. B. Overlaps of siRNA loci with NUMTs. Species in A and B annotated at the bottom. The numbers on top of each bar represent the enrichment, and the error bars represent the 95% confidence interval from random sampling of the genome.

## Conclusions

We introduce TIPPo, a user-friendly, reference-free approach for assembling plant organellar genomes. TIPPo provides a streamlined and universal assembly process without the need for external reference genomes. For both chloroplast and mitochondrial genomes, we provide assembly graphs. For chloroplast genomes, we provide in addition information on heteroplasmy. A limitation of our approach is that it can only use high-quality long reads, but we feel this is justified given that this technology underpins many of the ongoing large-scale genome sequencing and assembly projects (Rhie et al. 2021; Darwin Tree of Life Project Consortium 2022; Lewin et al. 2022). We also note that another newly released assembler for plant organellar genomes that comes from some of the colleagues leading these large-scale efforts is also restricted to the use of high-quality long reads (Zhou et al. 2024).

TIPPo outperforms all other tested assemblers for chloroplast genomes. Compared to chloroplast genomes, assessing the performance for mitochondrial genomes is more difficult due to the diversity of plant mitochondrial genomes. Based on the completeness of protein-coding genes, TIPPo outperforms the second-best tool PMAT (Bi et al. 2024) in eight species, while PMAT was superior for two species, *T. bicornis* and *H. pedunculosum*. A significant factor appears to be the presence of a large rDNA cluster in the nuclear genome of *T. bicornis*, which results in poor classification by Tiara (Karlicki et al. 2022), the initial tool used by TIPPo for selecting input reads for the assembly. The incomplete mitochondrial assembly of *H. pedunculosum* is likely the result of excessive filtering using a k-mer-based approach.

## Materials and Methods

### Data sources

HiFi datasets were downloaded from publicly available databases, details refer to **Table S1**. The accession numbers for chloroplast and mitochondrial genomes are provided in **Table S2** and **Table S3**. A phylogenetic tree of the 53 species was constructed with rtrees (https://github.com/daijiang/rtrees) (Li 2023).

### Evaluation of Tiara for read classification

First, minimap2 (2.24-r1122) with the parameter map-hifi was used to align all HiFi reads to the *A. thaliana* (Rabanal et al. 2022)and *O. sativa* (Shang et al. 2023)reference genomes, retaining only the primary alignments. Next, Tiara (1.0.3) (Karlicki et al. 2022) was used to classify HiFi reads as organellar. A 100 kb sliding window was applied to calculate the proportion of reads classified as organellar by Tiara compared to minimap2 in each window. The results were visualized using ggplot2 (3.5.1).

### Parameter selection for TIARA and Flye

For evaluating the impact of parameters on Flye, we tested: (1) default parameters; (2) default parameters with --meta; (3) default parameters with --keep-haplotypes; and (4) default parameters with both --meta and --keep-haplotypes (**Figure S18**). For selecting the best parameter for TIARA, we used different parameter combinations: k1 with 3 values (4, 5, 6), k2 with 4 values (4 to 7), and p with 15 values (0.3 to 1), resulting in a total of 180 combinations for reads classification.

### Assembly of organellar genomes

We used fxTools (v0.1.0) (https:/github.com/moold/fxTools) for subsampling PacBio HiFi reads to approximate 4x nuclear genome coverage for each species, except for 2x for *T. chinensis*, which has a particularly large nuclear genome (Xiong et al. 2021). For *S. conica*, with its large mitochondrial genome (Sloan et al. 2012), we used 10x nuclear genome coverage. For *Lycopodium japonicum*, the sequenced data coverage is only 0.59x (Bi et al. 2024). We used identical datasets for assembly with the different tools. TIPPo (v2.1) with default parameters was used to assemble chloroplast and mitochondrial genomes simultaneously. PMAT (v1.5.3) (Bi et al. 2024) is optimized for the assembly of plant mitochondrial genomes and has not been optimized for chloroplast assembly (**Figure S5**). For PMAT, the auto mode was firstly used with the parameters -tp mt and -tp all, applied separately. Subsequently, the buildgraph mode was applied using the output from the auto mode. For ptGAUL (v1.0.5) (Zhou et al. 2023) and CLAW (https:/github.com/aaronphillips7493/CLAW) (Phillips et al. 2024), which only assemble chloroplast genomes, the chloroplast genome sequences of closely related species were provided and run with default parameters.

### Whole-genome alignment and visualization

To compare genomes assembled from different sources, whole-genome alignments were performed with MiniTV (https://github.com/weigelworld/minitv), which uses minimap2 (v2.24-r1122) (Li 2018) for alignment, followed by visualization with AliTV (v1.0.6) (https:/alitvteam.github.io/AliTV/d3/AliTV.html) (Ankenbrand et al. 2017).

### Removal of chloroplast sequences from mitochondrial assemblies

First, we converted the mitochondrial assembly graphs into fragments. Given that the TIPPo chloroplast assembly results are the cleanest and complete, we aligned the mitochondrial contigs from PMAT (v1.5.3) (Bi et al. 2024) to the TIPPo chloroplast genome using minimap2 (2.24-r1122) (Li 2018). Contigs that were covered over more than 90% of their length by the chloroplast genome and had greater than 95% similarity to it were labeled as “chloroplast”. Using Bandage (v0.9.0) (Wick et al. 2015), we colored the nodes identified as chloroplast sequences in green and confirmed their identity after visual inspection. We removed the chloroplast sequences from the mitochondrial assemblies.

### Assessing assembly completeness

We obtained amino acid sequence files for 41 conserved mitochondrial genes from mitopy (https://dogma.ccbb.utexas.edu/mitofy/) (Alverson et al. 2010). We used BLASTX (2.9.0+) (McGinnis and Madden 2004) to align mitochondrial genome assemblies to each of the 41 genes, using a threshold of 1e-3. Considering that the current mitochondrial assembly results are presented in the format of an assembly graph, where long repeats will be collapsed into a single node, we evaluate gene completeness based on the presence or absence of genes, without accounting for their copy number.

### Performance benchmarking

All organellar genomes were assembled on an AMD EPYC 7742 processors with 64 cores and 1 TB of RAM. Runtime and peak memory usage were calculated using the /usr/bin/time -v command. All the assembly tools were set to run with 40 threads.

### NUMT and NUPT analysis

To identify NUPTs and NUMTs in the nuclear genome, we used BLASTN (2.9.0+) (McGinnis and Madden 2004) with the parameters -evalue 1e-5, -dust no, -penalty -2, -word_size 9, -outfmt 6. We aligned the chloroplast and mitochondrial genomes to their respective nuclear genomes and retained hits with an identity of > 80% and a length > 100 bp. Considering the redundancy in the BLASTN output, we removed all high-scoring segment pairs (HSPs) completely embedded in longer HSPs. We merged overlapping HSPs with bedtools (v2.31.1) (Quinlan and Hall 2010). The identity of the merged interval in the nuclear genome to the organellar genome was calculated as the average of the identities before merging.

To identify substitutions in NUPTs and NUMTs relative to the chloroplast and mitochondrial genomes, we used minimap2 (version 2.24-r1122) (Li 2018) with the parameters --paf-no-hit -ax asm5 --cs -r2k to generate alignment files. Finally, we used htsbox (version r345) (https://github.com/lh3/htsbox) with the parameters pileup -q5 -evcf to call variants.

### Annotation of siRNA loci and overlap with NUPTs/NUMTs

For each of the selected 11 species, we downloaded data from two libraries. We used ShortStack (v4.0.4) (Axtell 2013) with default parameters to annotate siRNA loci. In short, reads with one or no mismatch were retained, and multi-mapping reads were assigned to a single location with the U model. GAT (v1.3.5) (Heger et al. 2013) was used to test whether the siRNA locus overlaps were greater than expected by chance with the parameter -num-samples=1000.

## Data availability

Chloroplast and mitochondrial assembly graphs are available on Figshare at https://doi.org/10.6084/m9.figshare.26362141.v1.

## Code availability

TIPPo is available at Github (https://github.com/Wenfei-Xian/TIPP). Code to reproduce results from this paper can be found at Github (https://github.com/Wenfei-Xian/Reproducible_for_TIPP_paper).

## Acknowledgements

We thank Andrea Movilli, Adrian Contreras, Yueqi Tao, Svitlana Sushko, Li He and Haim Ashkenazy for the discussions. This work was supported by the Max Planck Society and the Novozymes Prize of the Novo Nordisk Foundation (DW).

## Author Contributions

W.X. designed the project and conducted the analyses. I.B. set up the computational environment. I.B., Z.B., S.V. and A.G. tested the tool. D.W. supervised the project. W.X. wrote the first draft of the manuscript. W.X. and D.W. prepared the final manuscript with input from all authors.

