## Supplementary Figures for "TIPPo: A User-Friendly Tool for De Novo Assembly of Organellar Genomes with HiFi Data"

*Arabidopsis thaliana* - Col-0

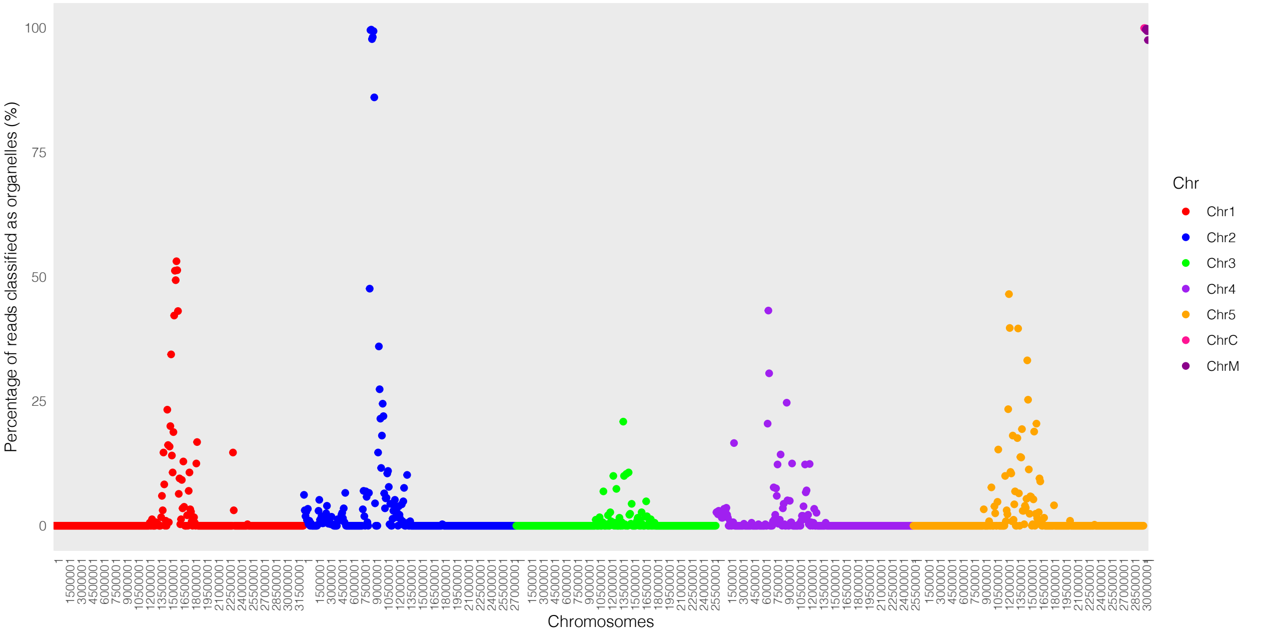


*Oryza sativa* ssp. japonica cv. Nipponbare

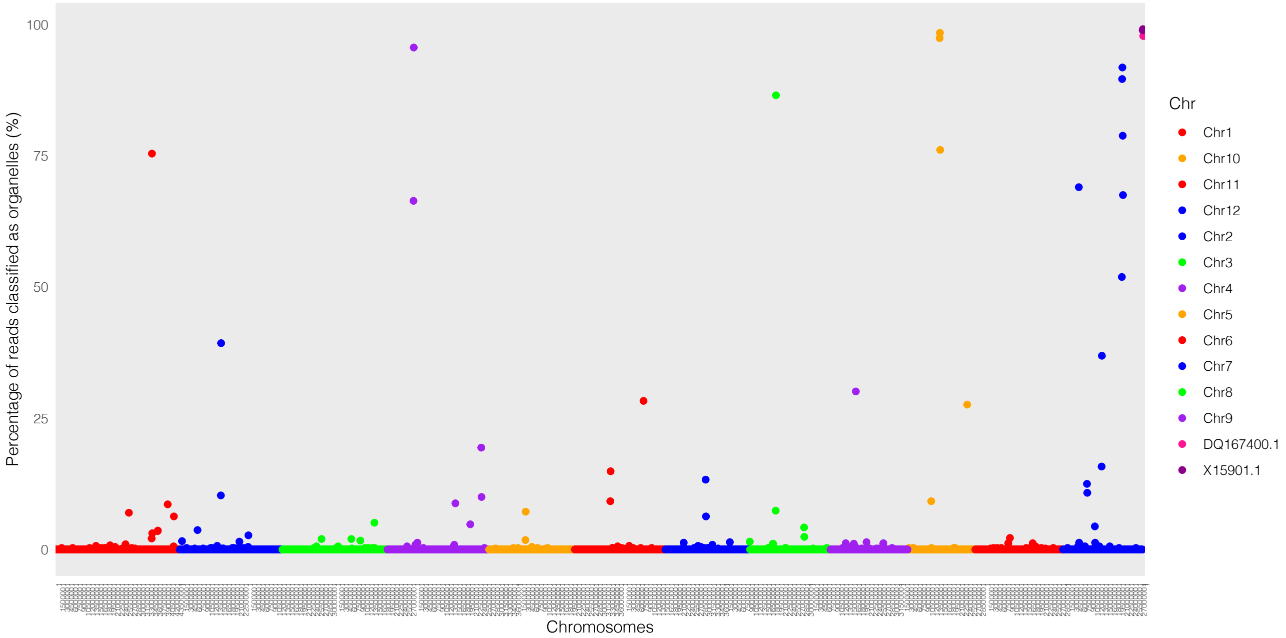


Figure S1. Testing the performance of TIARA in two plants.

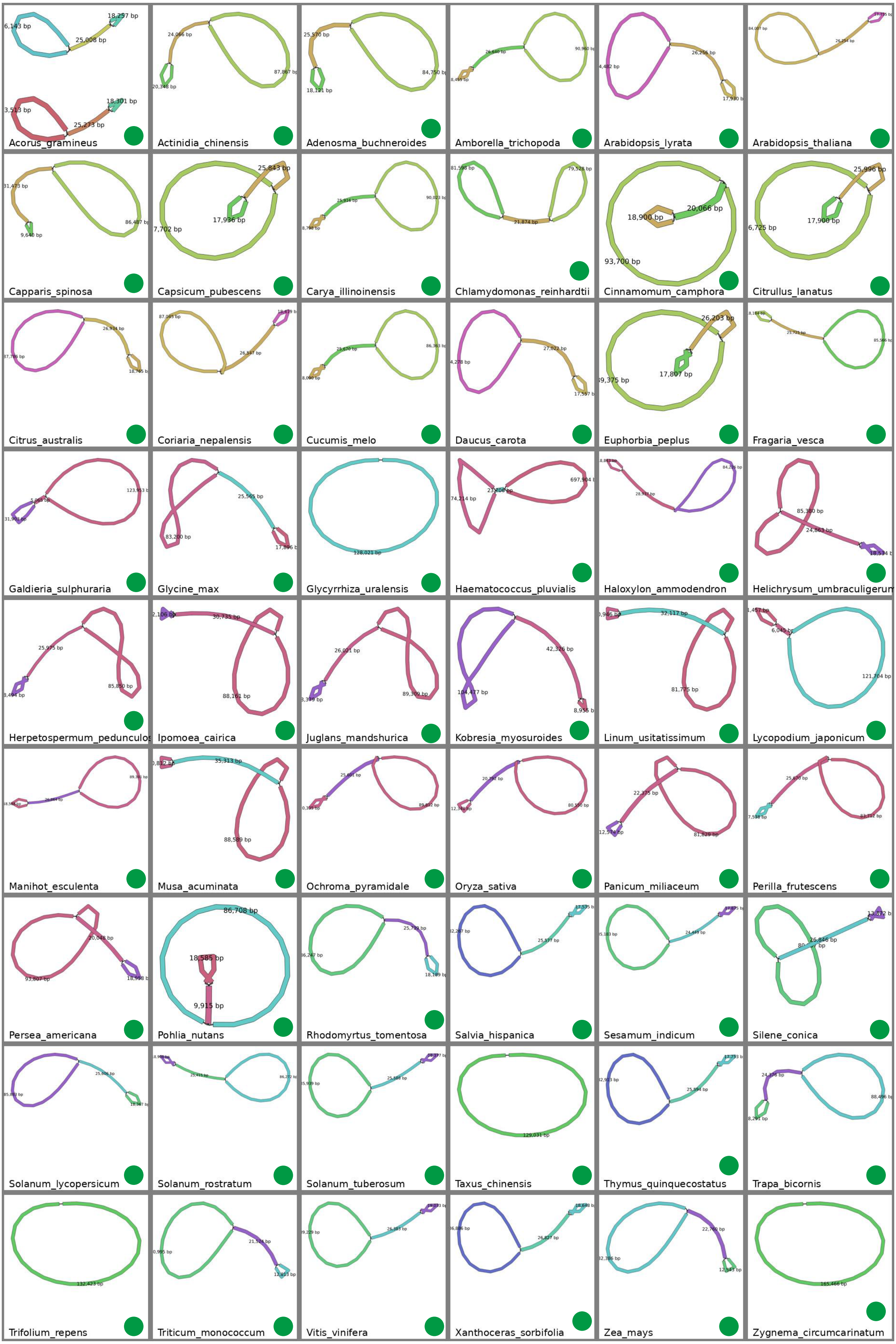


Figure S2. Chloroplast genomes of 54 species assembled using TIPPo. Green solid dots represent class 1 complete genomes. Blue solid dots represent class 2 complete genomes and other sequences. Brown solid dots represent class 3 incomplete assemblies.

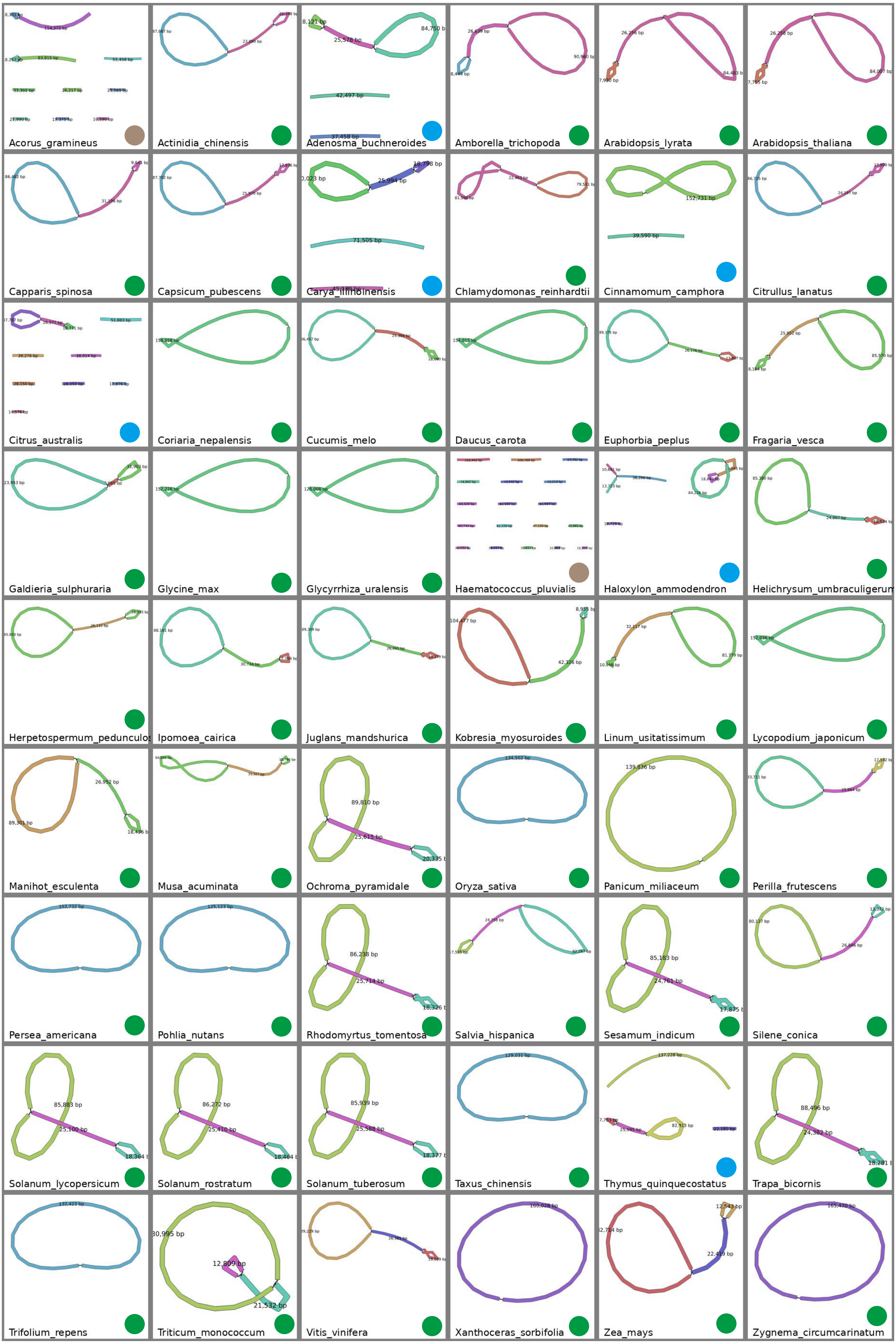


Figure S3. Chloroplast genomes of 54 species assembled using ptgaul. Green solid dots represent class 1 complete genomes. Blue solid dots represent class 2 complete genomes and other sequences. Brown solid dots represent class 3 incomplete assemblies.

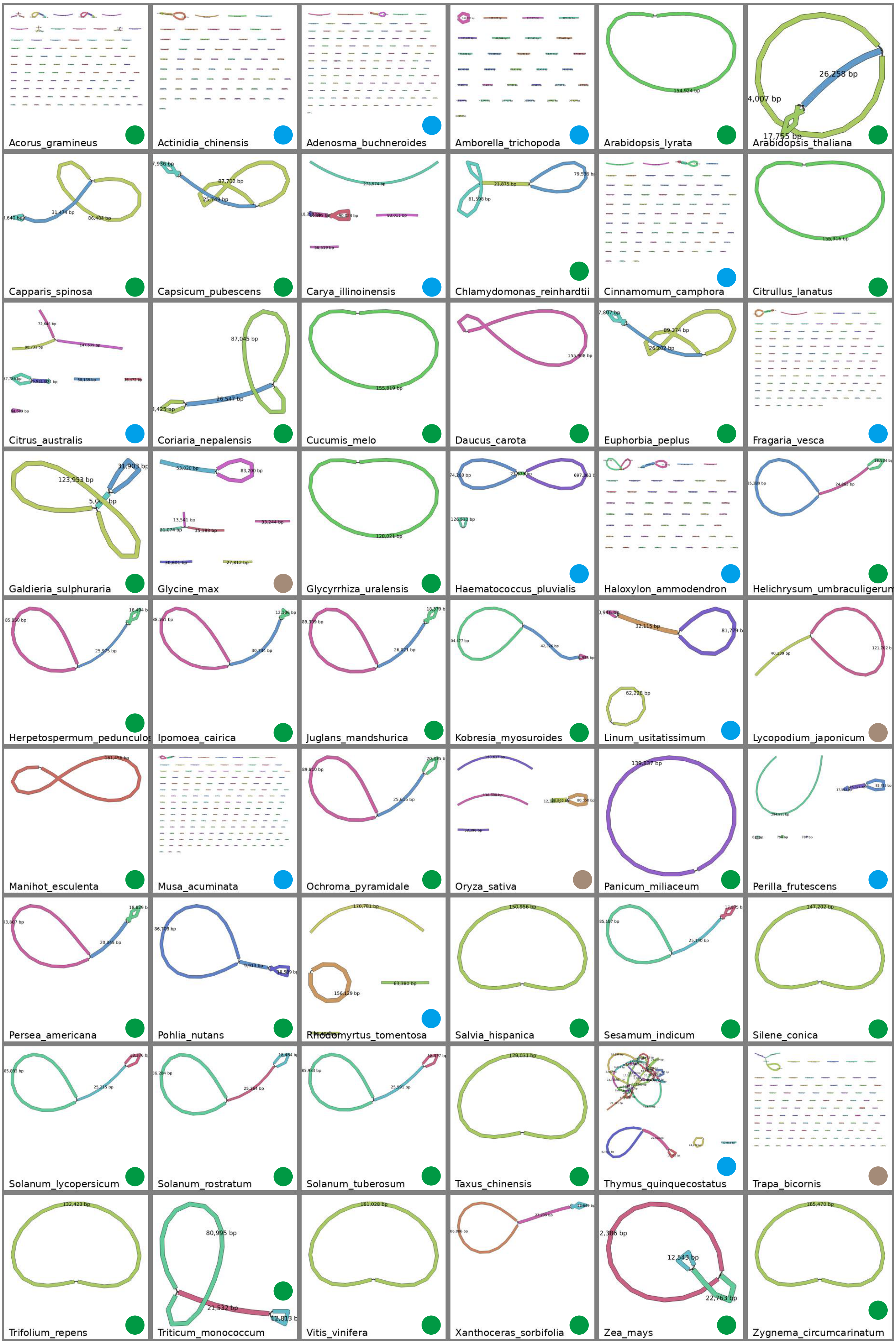


Figure S4. Chloroplast genomes of 54 species assembled using CLAW. Green solid dots represent class 1 complete genomes. Blue solid dots represent class 2 complete genomes and other sequences. Brown solid dots represent class 3 incomplete assemblies.

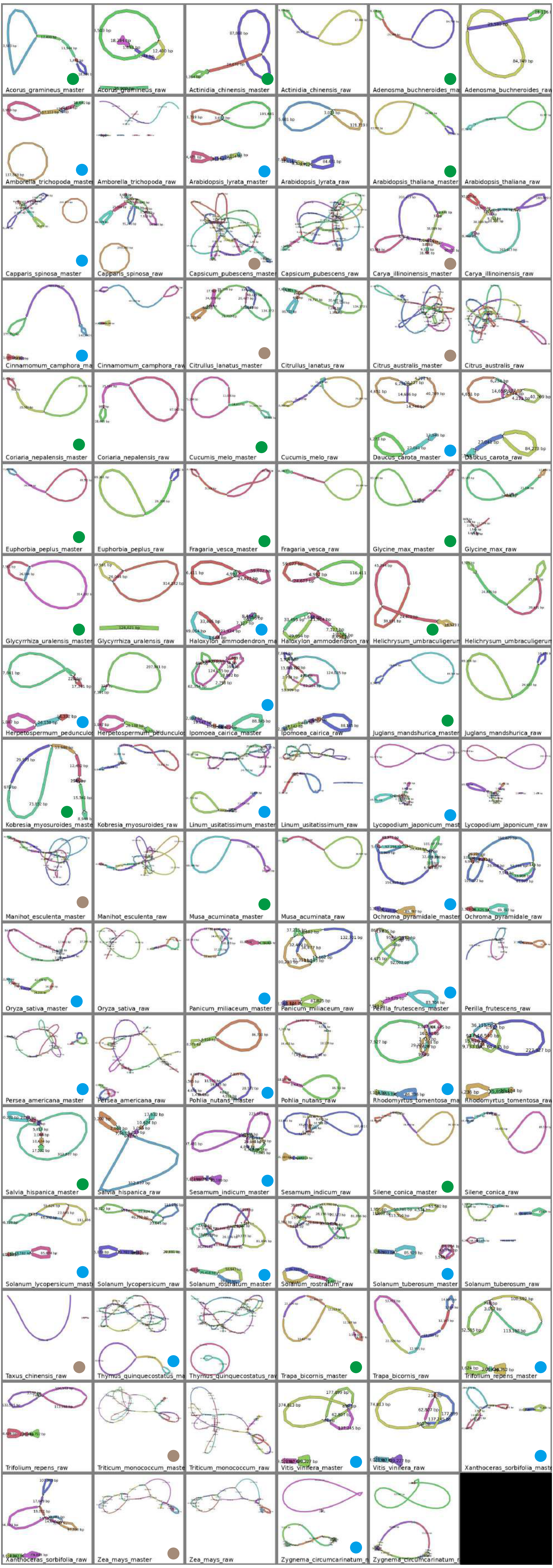


Figure S5. Chloroplast genomes of 54 species assembled using PMAT pt. Green solid dots represent class 1 complete genomes. Blue solid dots represent class 2 complete genomes and other sequences. Brown solid dots represent class 3 incomplete assemblies.

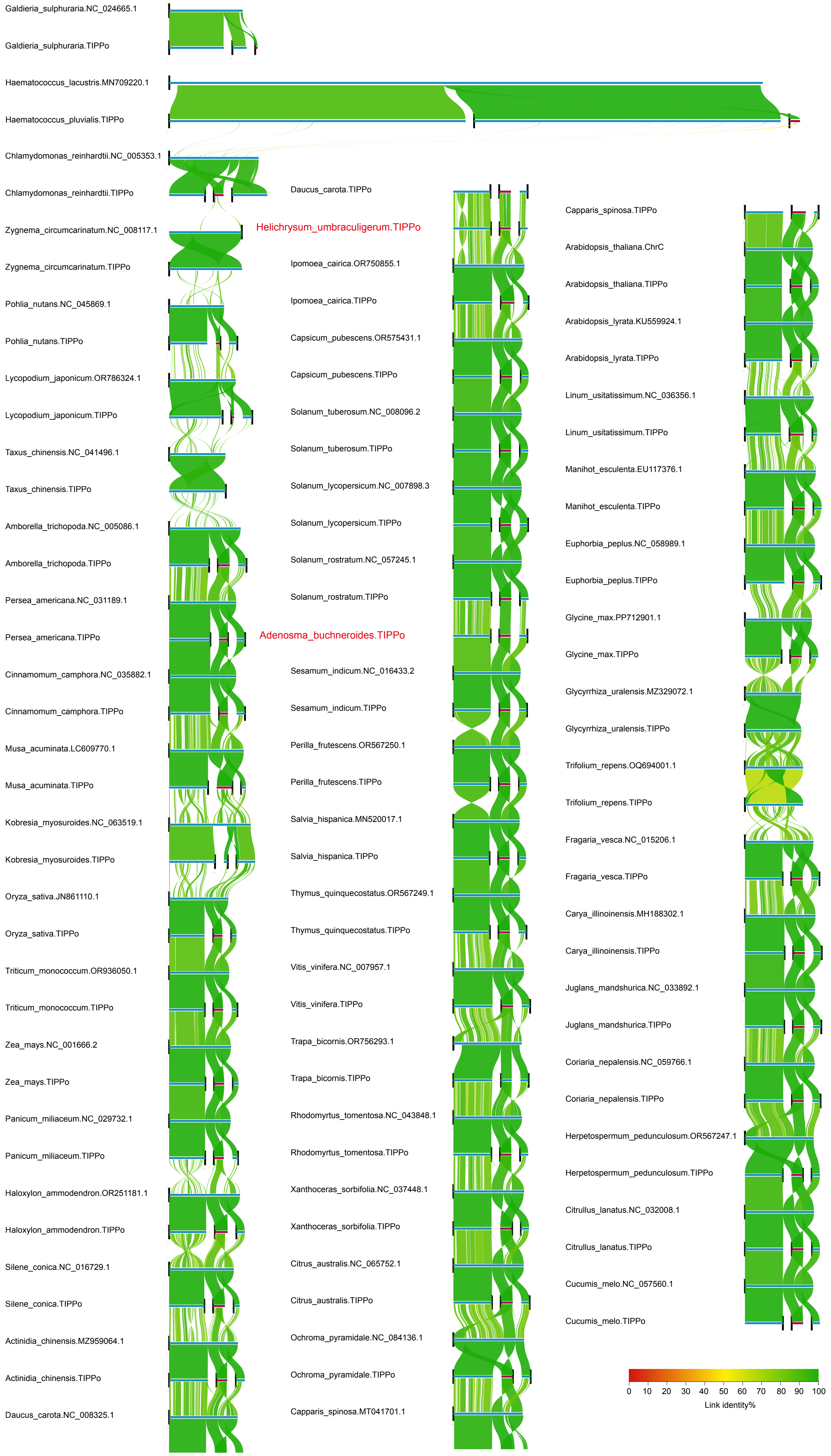


Figure S6. Whole genome alignment of chloroplast genomes. Species names ending with "TIPPo" were assembled using the TIPPo tool, while those ending with an accession ID were downloaded from NCBI.

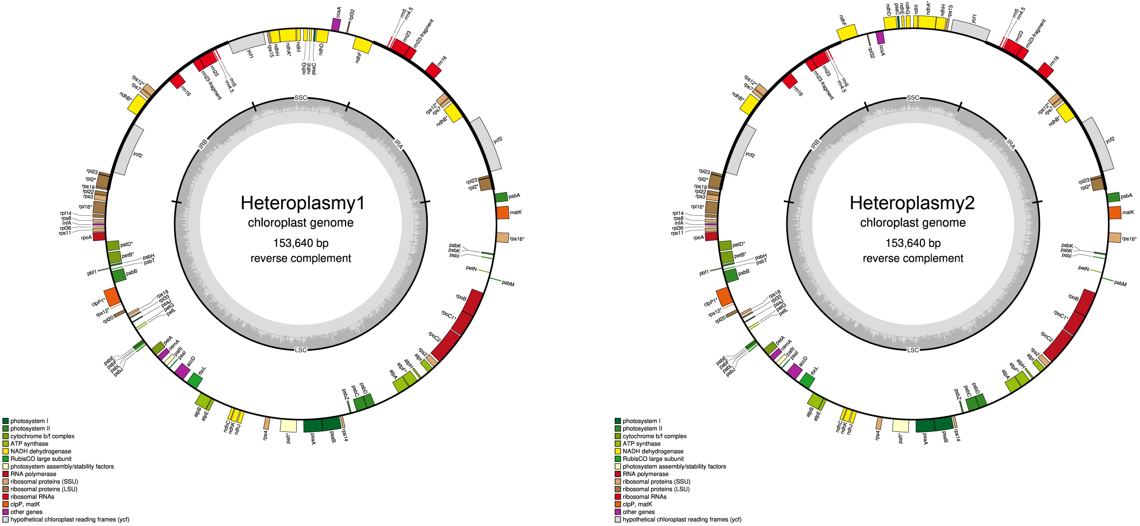


Figure S7. Chloroplast genome of *Adenosma buchneroides.*

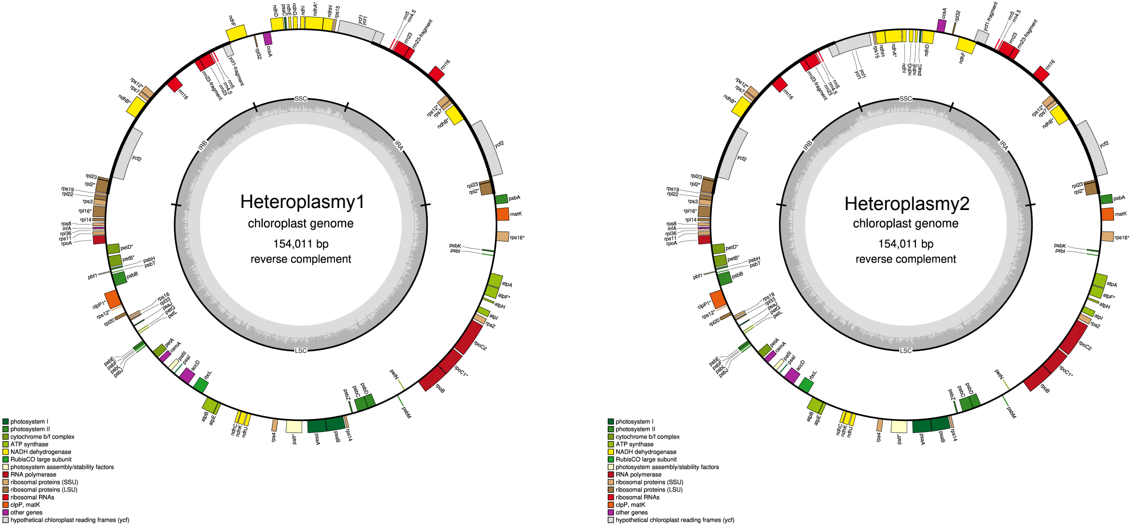


Figure S8. Chloroplast genome of *Helichrysum umbraculigerum.*

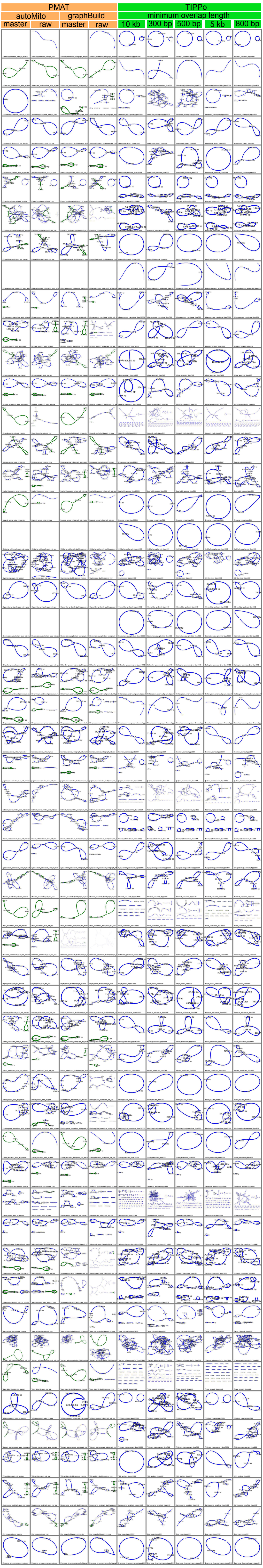


Figure S9: Mitochondrial assembly graph of PMAT (auto-raw, auto-master, graphbuild-raw, graphbuild-master) and TIPPo (minimum overlap length: 300, 500, 800, 5000 and 1000 bp). Green nodes represent chloroplast sequences, and blue nodes represent non-chloroplast sequences.

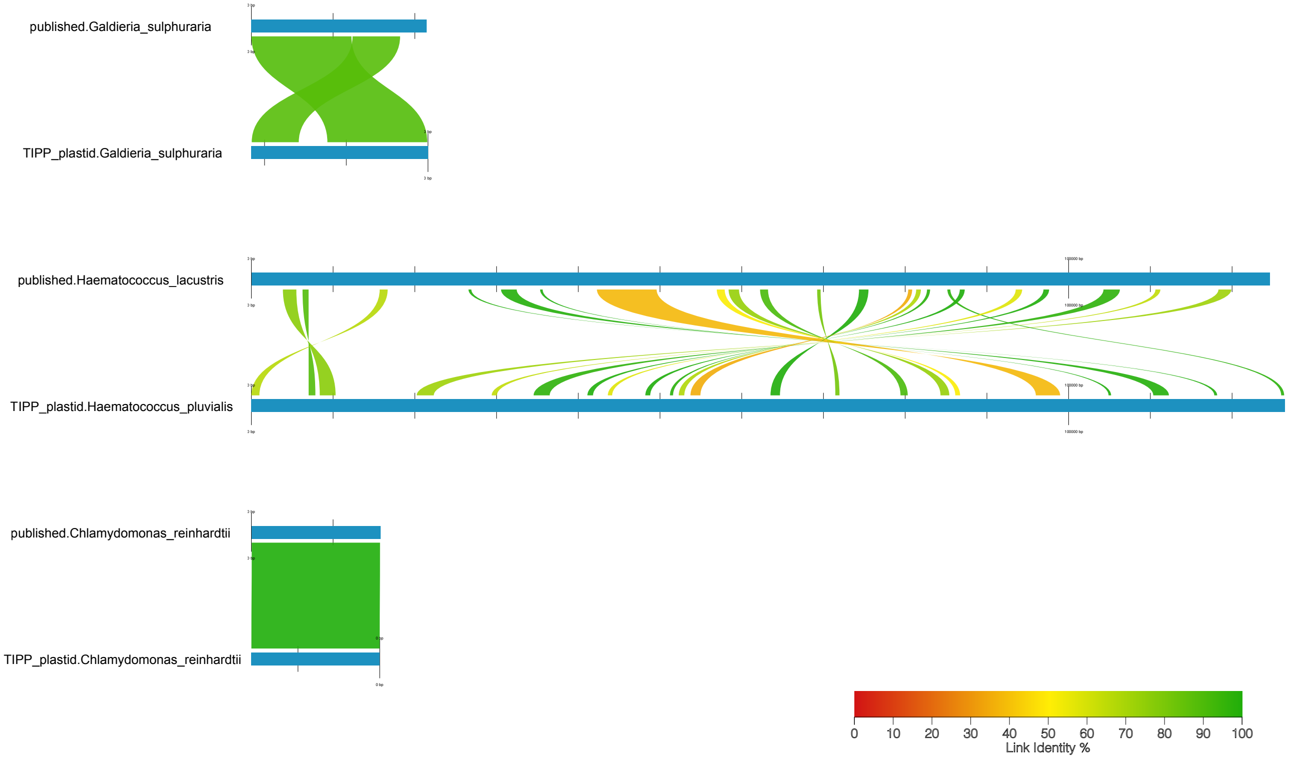


Figure S10. Whole genome alignment of mitochondrial genomes.

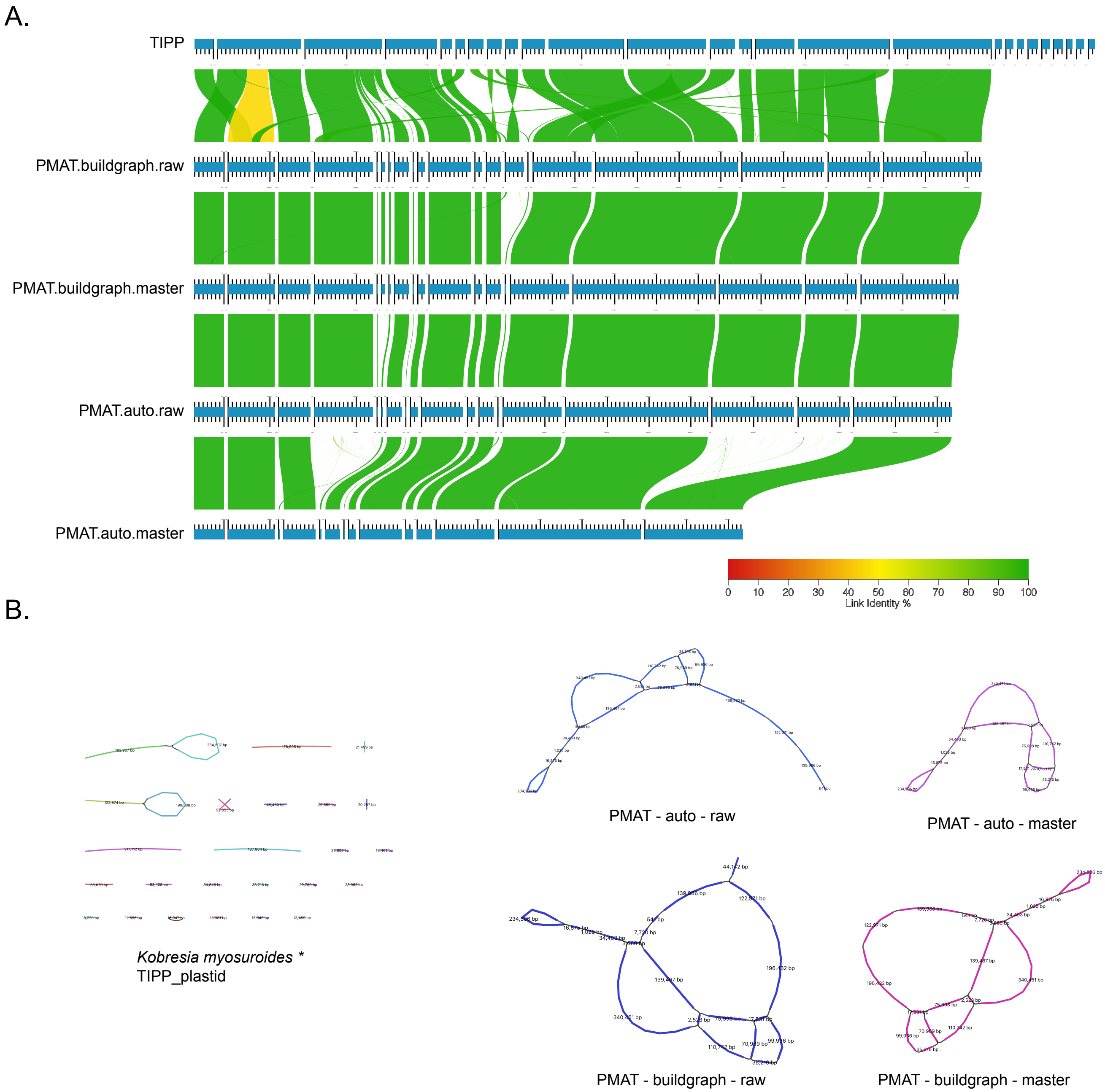


Figure S11. Mitchondrial assemblies of *Kobresia myosuroides*. A. whole genome alignment. B. visualization of assembly graphs.

A.

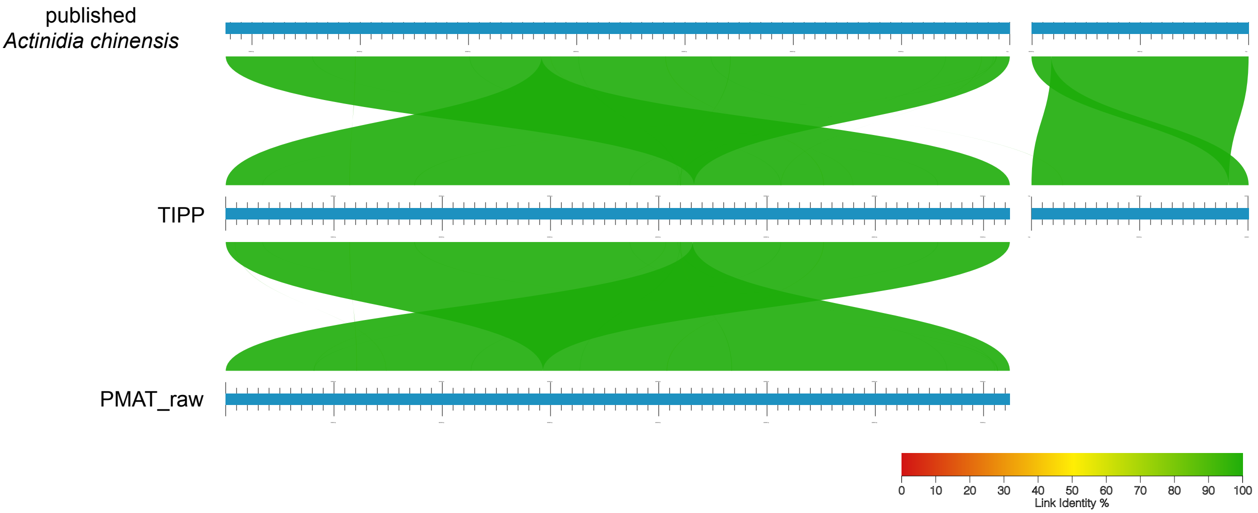


B.

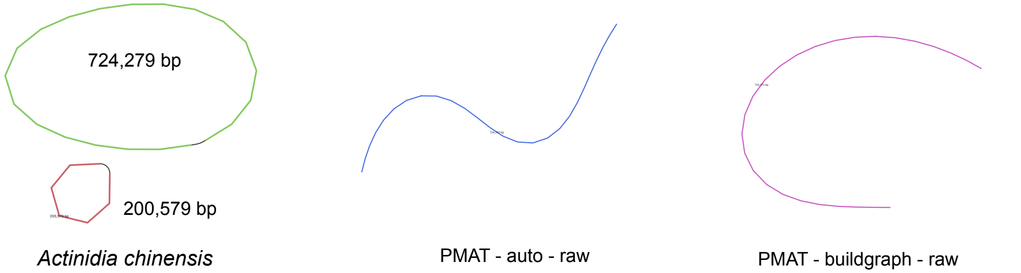


No difference between auto mode and buildgraph mode.

C.

```
/tmp/global2/wxian/software/PMAT/bin/PMAT graphBuild -rs Actinidia_chinensis.4X.fastq.gz -o Actinidia_chinensis.PMAT_buildGraph  -gs 616M -c Actinidia_chinensis.PMAT/assembly_result/PMATContigGraph.txt
 -a Actinidia_chinensis.PMAT/assembly_result/PMATAllContigs.fna
Reading files ...
2024-09-18 11:02:21
[ INFO 2024-09-18 11:02:25 ] Contig number : 65087
[ INFO 2024-09-18 11:02:25 ] Longest Contig : contig1 724272bp
[ INFO 2024-09-18 11:02:25 ] ------------------------------------------------------------
Candidate seeds search start ...
2024-09-18 11:02:25

BLASTn encountered an error:
Warning: [blastn] Examining 5 or more matches is recommended

[ INFO 2024-09-18 11:04:24 ] 5 contigs are used as candidate seeds
Contigs         Length          Depth
________________________________________________
contig00001     724272bp        27.8X
contig19935     22465bp         4.0X
contig03346     137721bp        3.5X
contig37452     24411bp         2.7X
contig49476     10081bp         1.7X
[ INFO 2024-09-18 11:04:25 ] ------------------------------------------------------------
Seeds extension start ...
2024-09-18 11:04:25
Mt Extension No.1: 100% |####################################################################################################################
##########################|
Seeds extension end ...
2024-09-18 11:04:25
[ INFO 2024-09-18 11:04:25 ] ------------------------------------------------------------
[ INFO 2024-09-18 11:04:32 ] Start generating the gfa file ...
[ INFO 2024-09-18 11:04:57 ] save gfa for 23.73s
[ INFO 2024-09-18 11:04:57 ] 1 contigs are added to a raw graph
[ ERROR 2024-09-18 11:04:57 ] There is no master structure for this seeds extension result.
[ INFO 2024-09-18 11:04:57 ] Generate gfa task end.
[ INFO 2024-09-18 11:04:57 ] Task over, bye!
```

Figure S12. Mitchondrial genome assemblies of *Actinidia chinensis*.
A. whole genome alignment. B. visualization of assembly graphs. C. log of PMAT buildGraph.

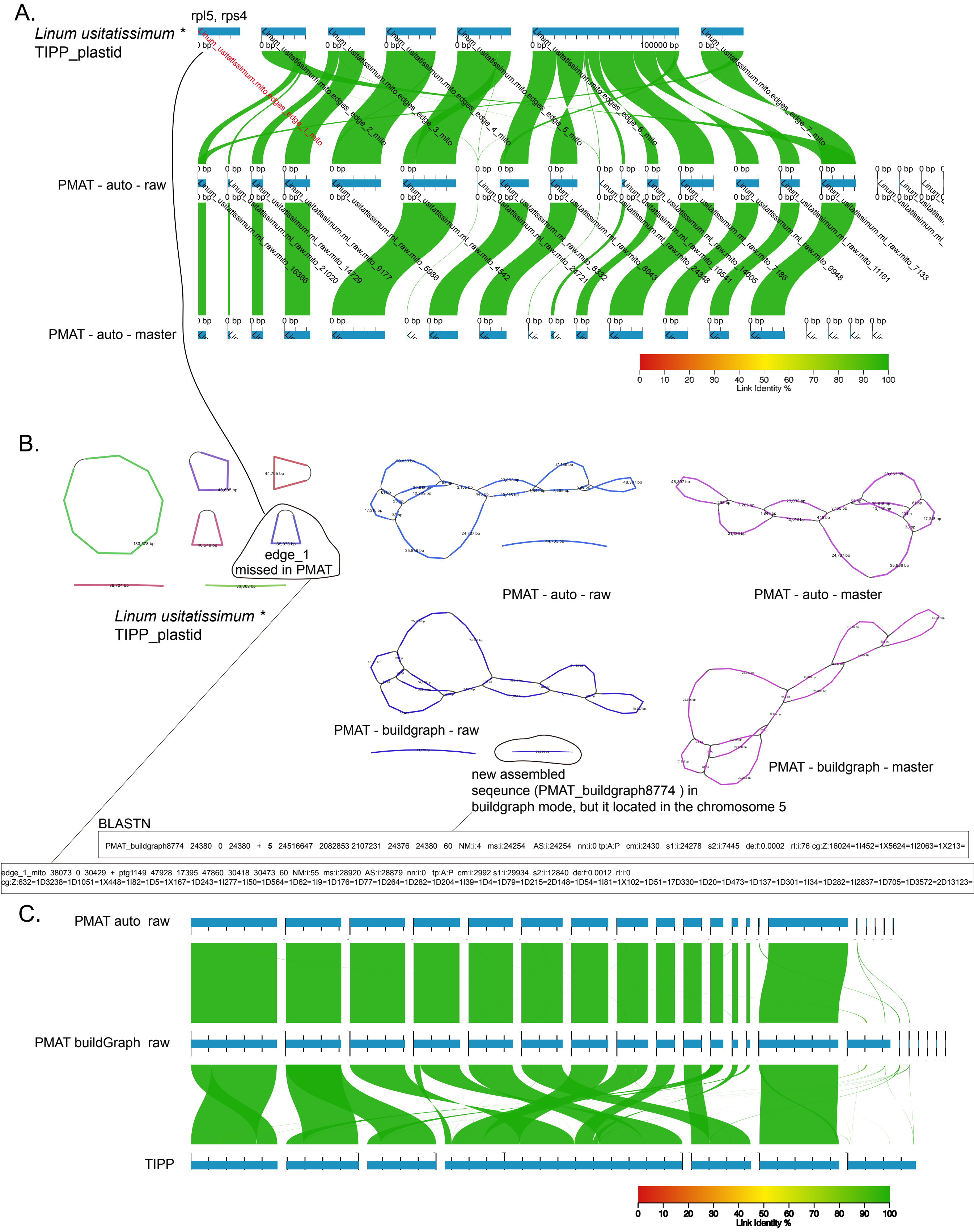


Figure S13. Mitchondrial genome assemblies of *Linum usitatissimum*.
A. whole genome alignment of TIPPo, PMAT-auto_raw and PMAT-auto-master assemblies.
B. visualization of assembly graphs. C. Whole genome alignment of TIPPo, PMAT-auto-raw and PMAT-buildgraph-raw.

A.

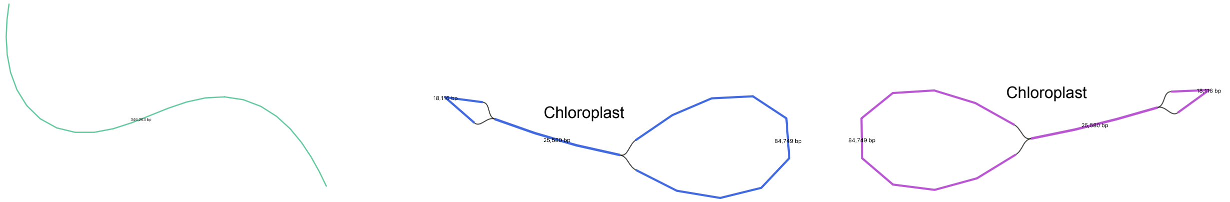


*Adenosma buchneroides* *
TIPP_plastid

PMAT - raw

PMAT - master

In buildGraph model, only chloroplast genome is presented.
No difference between auto model and buildgraph model.

B.

```
/tmp/global2/wxian/software/PMAT/bin/PMAT graphBuild -rs Adenosma_buchneroides.4X.fastq.gz -o Adenosma_buchneroides.PMAT_buildGraph  -gs 442M -c Adenosma_buchneroides.PMAT/assembly_result/
PMATContigGraph.txt -a Adenosma_buchneroides.PMAT/assembly_result/PMATAllContigs.fna
Reading files ...
2024-09-18 11:33:05
[ INFO 2024-09-18 11:33:05 ] Contig number : 5691
[ INFO 2024-09-18 11:33:05 ] Longest Contig : contig1 1637434bp
[ INFO 2024-09-18 11:33:05 ] ------------------------------------------------------------
Candidate seeds search start ...
2024-09-18 11:33:05

BLASTn encountered an error:
Warning: [blastn] Examining 5 or more matches is recommended

[ INFO 2024-09-18 11:34:12 ] 5 contigs are used as candidate seeds
Contigs         Length          Depth

contig02570     25580bp         200.5X
contig00118     512004bp        4.4X
contig02179     19298bp         2.8X
contig03105     14421bp         1.0X
contig03809     6545bp          1.0X
[ INFO 2024-09-18 11:34:13 ] ------------------------------------------------------------
Seeds extension start ...
2024-09-18 11:34:13
Mt Extension No.1: 100% |####################################################################################################################################################
########################|
Mt Extension No.2: 100% |####################################################################################################################################################
########################|
Seeds extension end ...
2024-09-18 11:34:13
[ INFO 2024-09-18 11:34:13 ] ------------------------------------------------------------
[ INFO 2024-09-18 11:34:16 ] Start generating the gfa file ...
[ INFO 2024-09-18 11:34:32 ] save gfa for 14.83s
[ INFO 2024-09-18 11:34:32 ] 3 contigs are added to a raw graph
[ INFO 2024-09-18 11:34:32 ] 3 contigs are added to a master graph
[ INFO 2024-09-18 11:34:32 ] Generate gfa task end.
[ INFO 2024-09-18 11:34:32 ] Task over, bye!
```

Figure S14. Mitchondrial genome assemblies of *Adenosma buchneroides*.
A. Visualization of assembly graphs. B. log of PMAT buildGraph.

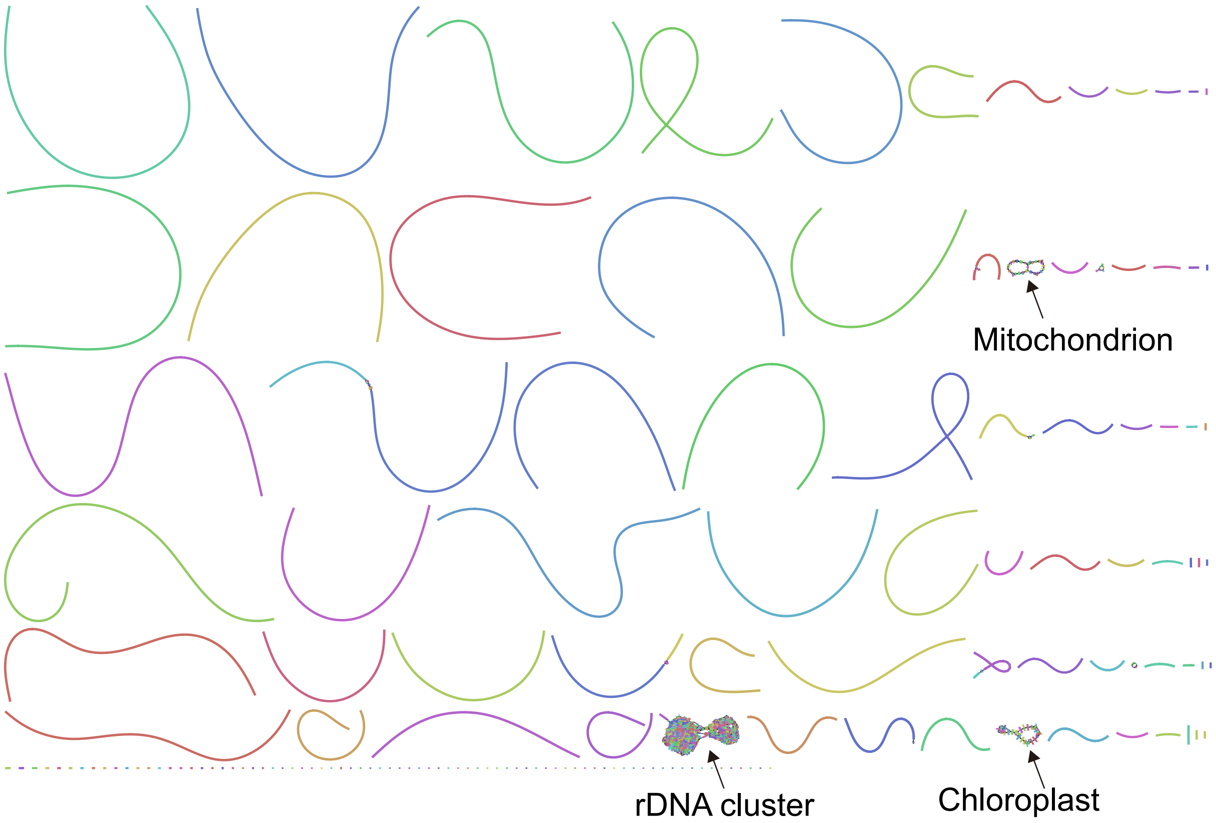


Figure S15. visualization of whole genome assembly graph of *Trapa bicornis.*

A.

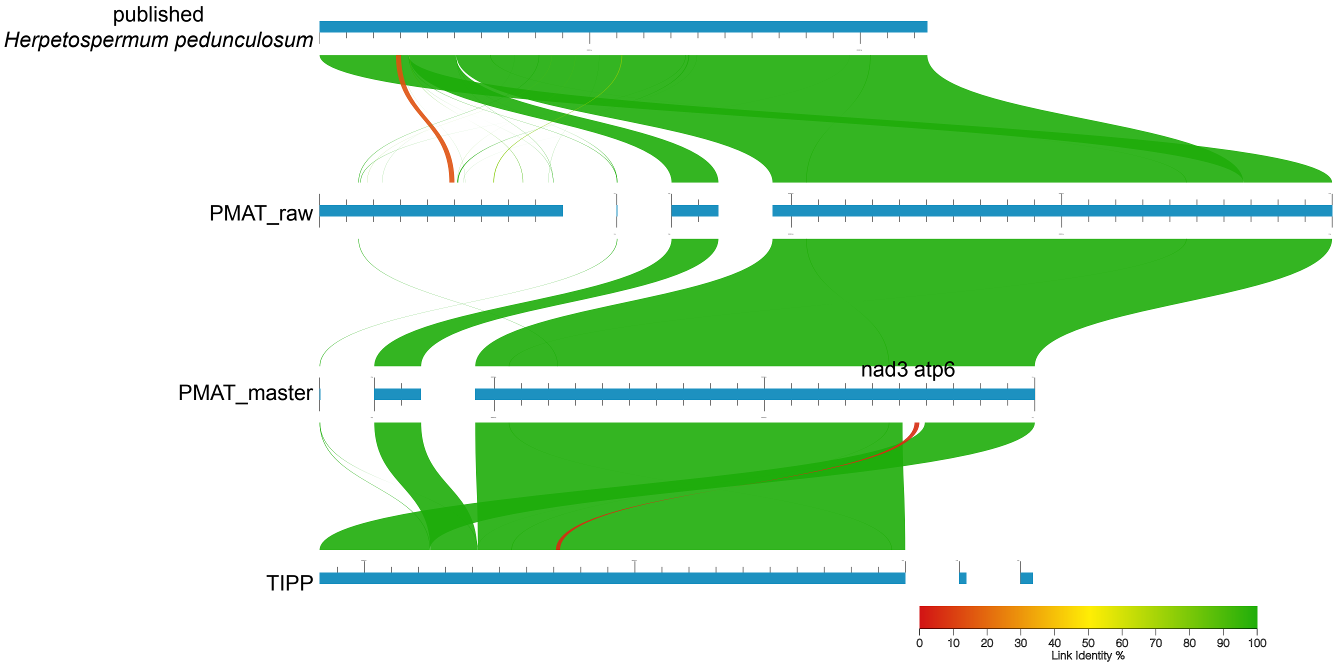


B.

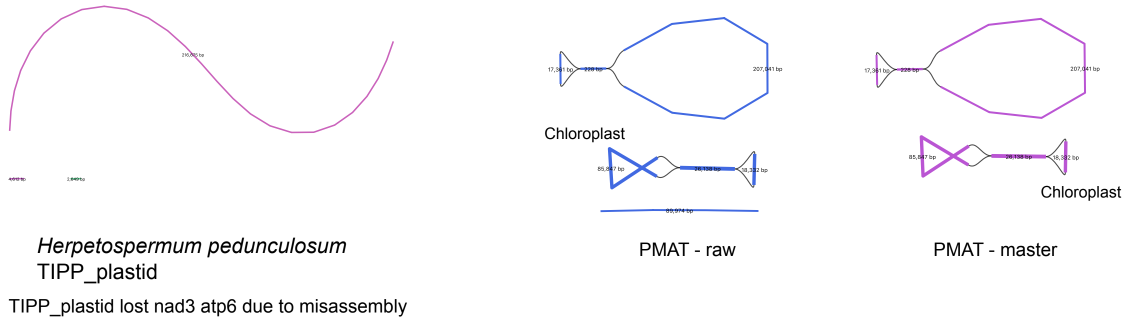


Figure S16. Mitchondrial genome assemblies of *Herpetospermum pedunculosum*. A. whole genome alignment. B. visualization of assembly graphs.

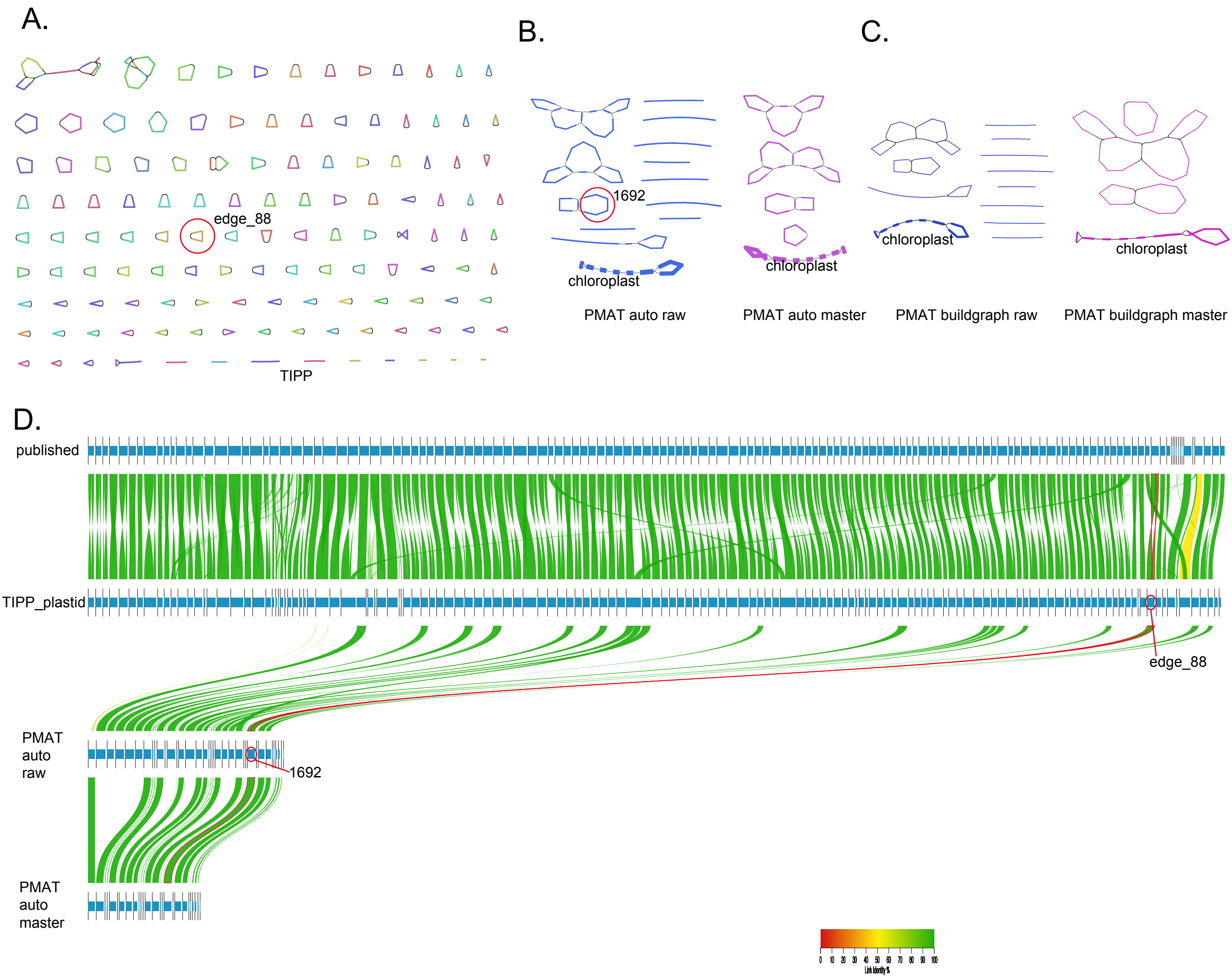


Figure S17. Assembly graph of *Silene conica.* A. assembly graph of TIPPo. B, assembly graph of PMAT auto model. C, assembly graph of PMAT buildgraph model. D. Whole genome alignment of published, TIPPo assembly, PMAT auto raw and PMAT auto master assembly. Blue bars represent the nodes in the assembly graph. Such as the node1692 in PMAT auto raw assembly show high similarity with the node edge_88 in TIPPo.

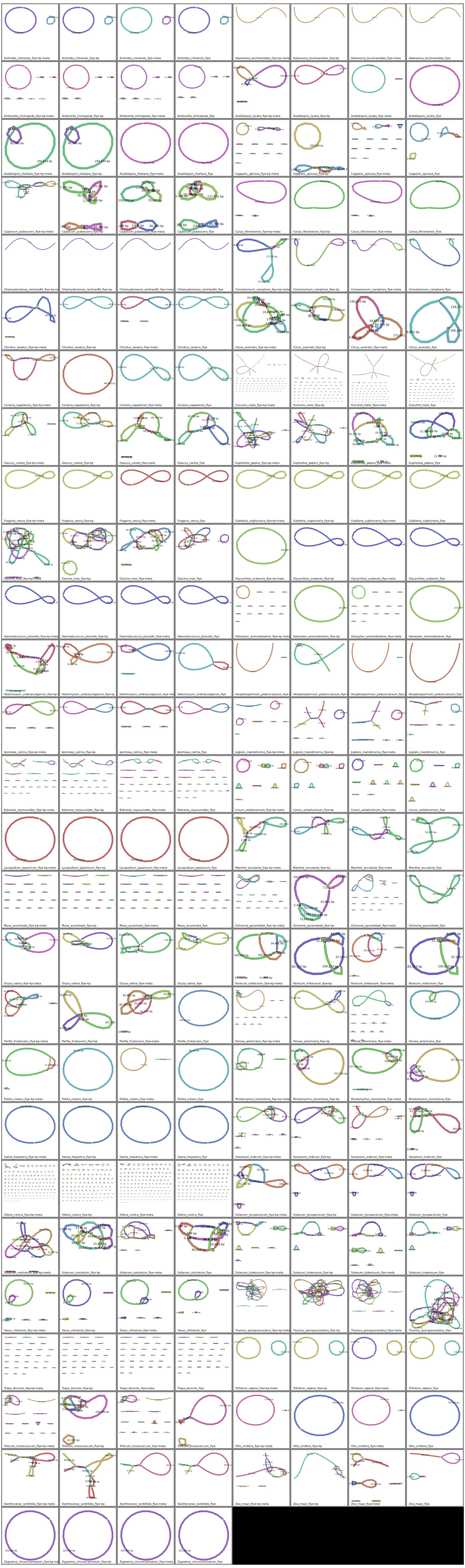

Figure S18: Mitochondrial assembly graph of TIPPo with different Flye parameters.
Flye default (species_flye), Flye --meta (species_flye-meta),
Flye --keep-haplotype (species_flye-kp), Flye --meta --keep-haplotype (species_flye-kp-meta).

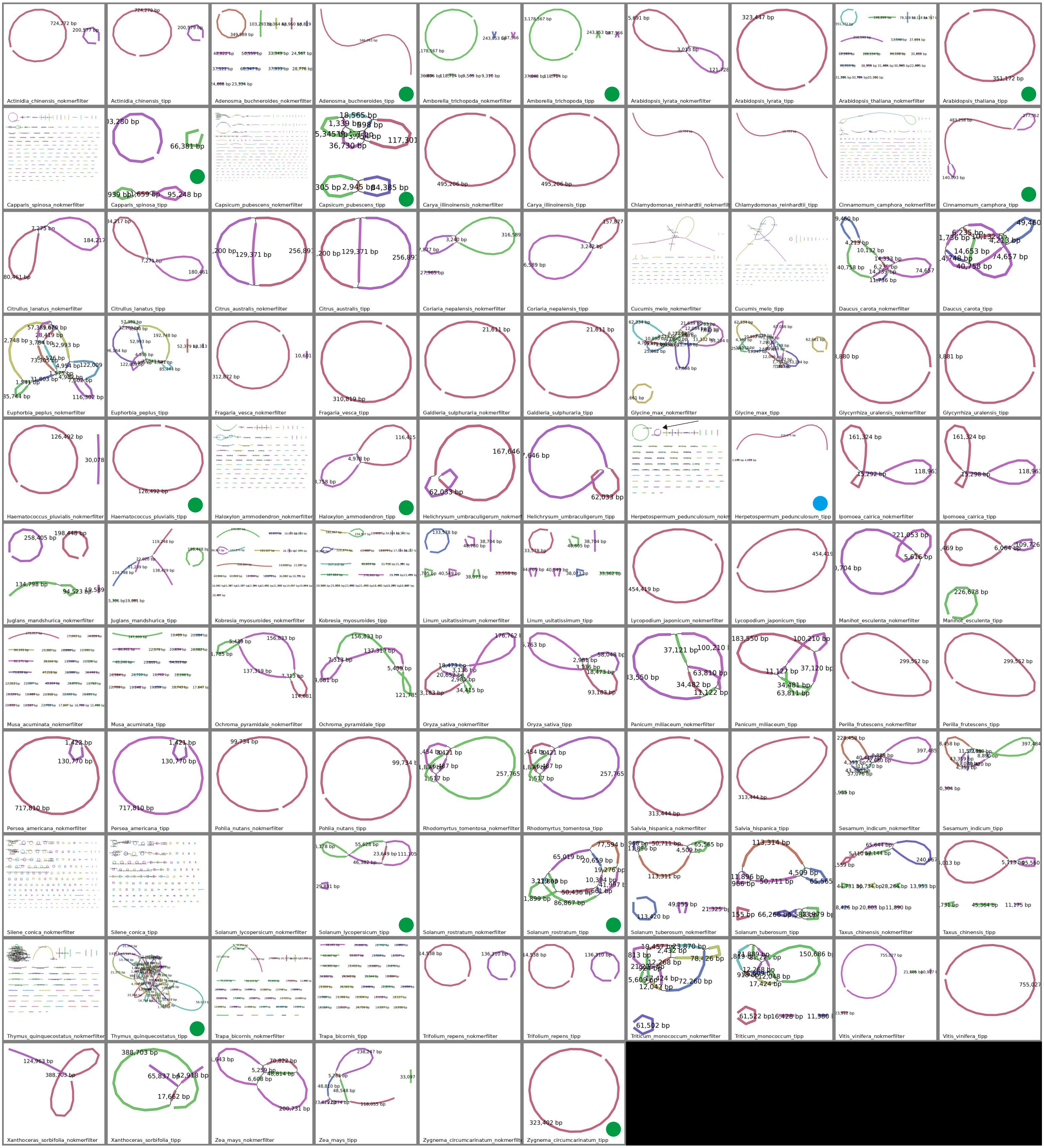


Figure S19: Mitochondrial assembly graph of TIPPo (species_tipp) and TIARA + Flye (species_nokmerfilter). Green solid dots represent that assemblies benefited from k-mer filtering. Blue solid dot represent that overfiltering by k-mer filtering.

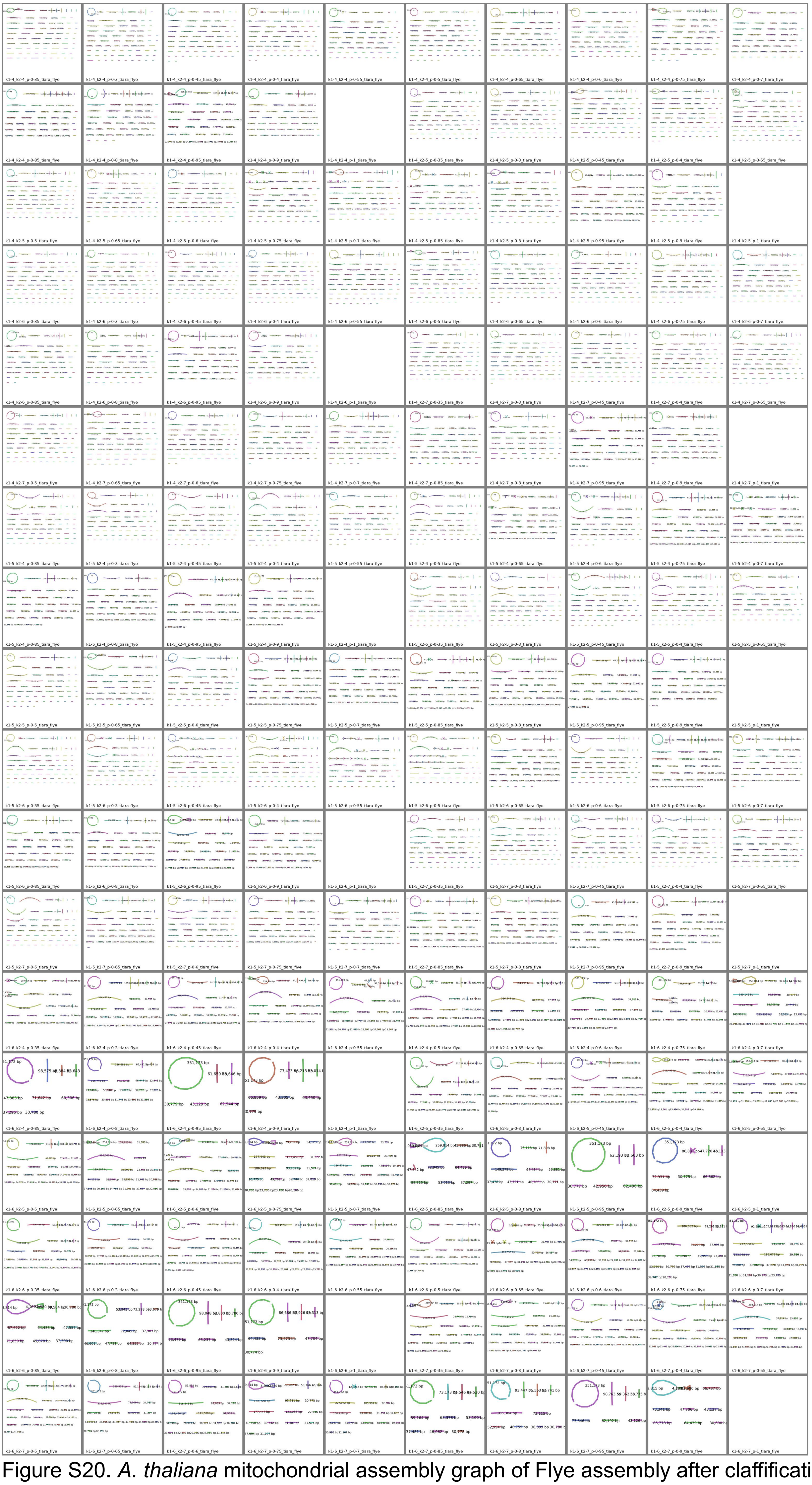

Figure S20. *A. thaliana* mitochondrial assembly graph of Flye assembly after claffification with TIARA with different parameter combinations: k1 with 3 values (4, 5, 6), k2 with 4 values (4 to 7), and p with 15 values (0.3 to 1), resulting in a total of 180 combinations.

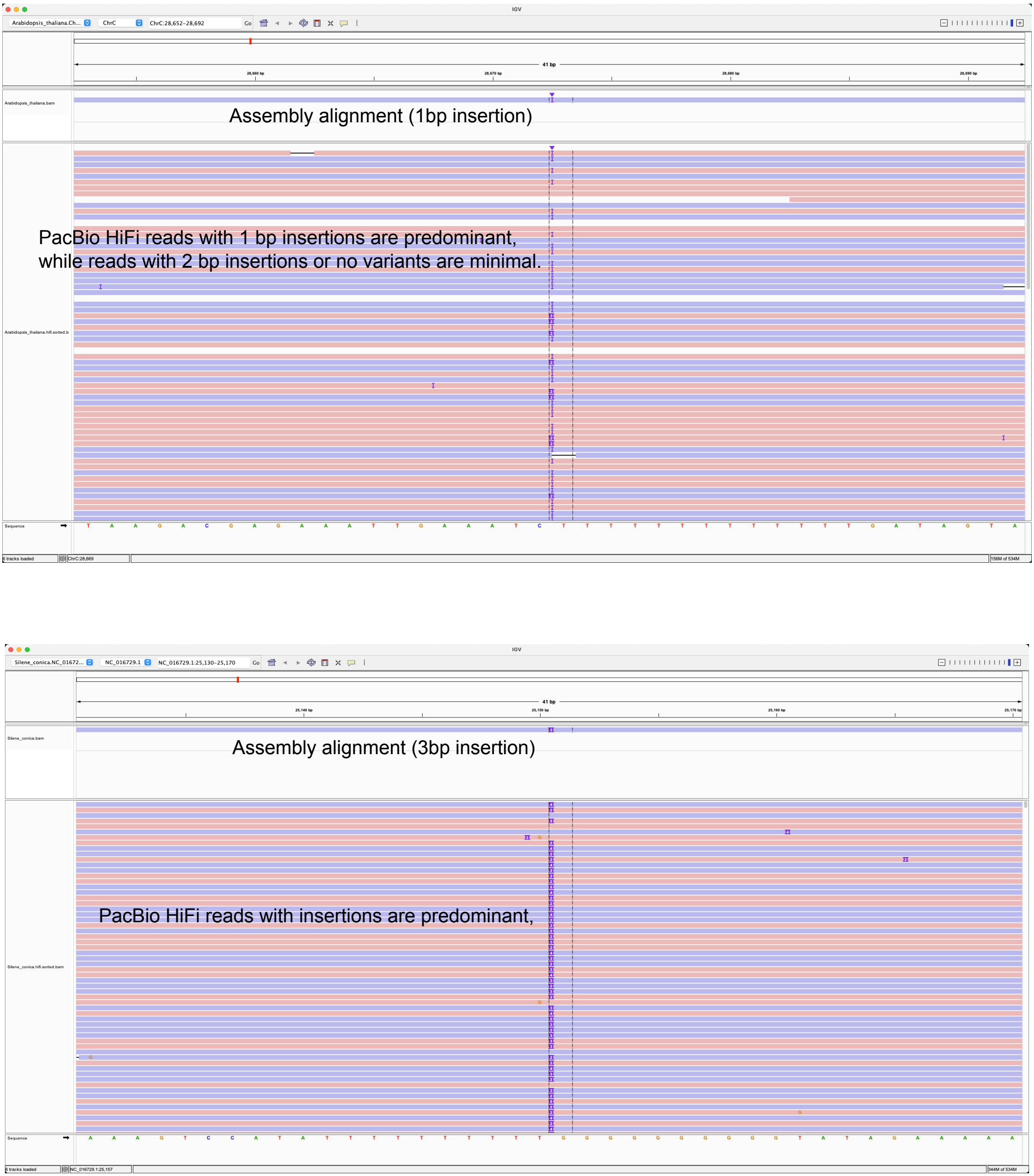


Figure S21. Base consensus compared with the published genomes. A. *Arabidopsis thaliana* Col-0 chloroplast. B. *Silene conica* chloroplast.

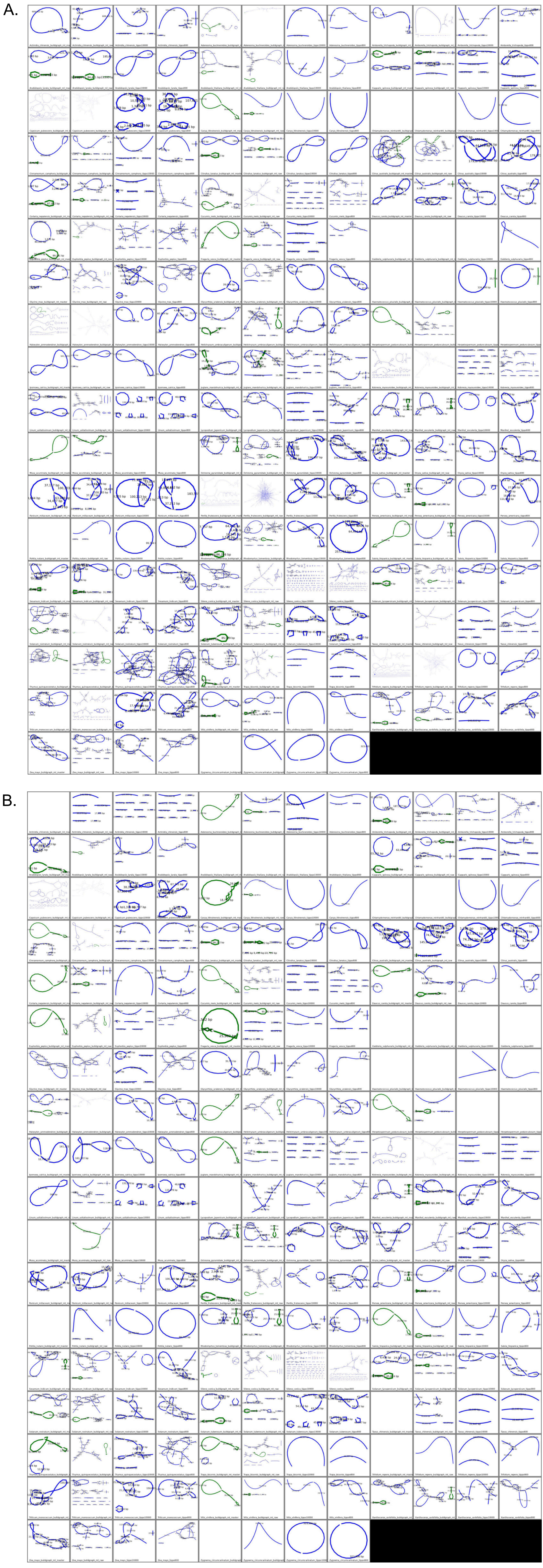

Figure S22: Mitochondrial assembly graph of PMAT (buildgraph-raw, buildgraph-master) and TIPPo (minimum overlap length: 800 and 10000 bp). A. mitochondrial assembly using 1x coverage for each species, except for 0.5x for *T. chinensis*, 2.5x for *S. conica* and 0.15x for *L. japonicum*. B. mitochondrial assembly using 0.5x for each species, except for 0.25x for *T. chinensis*, 1.25x for *S. conica* and 0.075x for *L. japonicum*. Green nodes represent chloroplast sequences, and blue nodes represent non-chloroplast sequences.

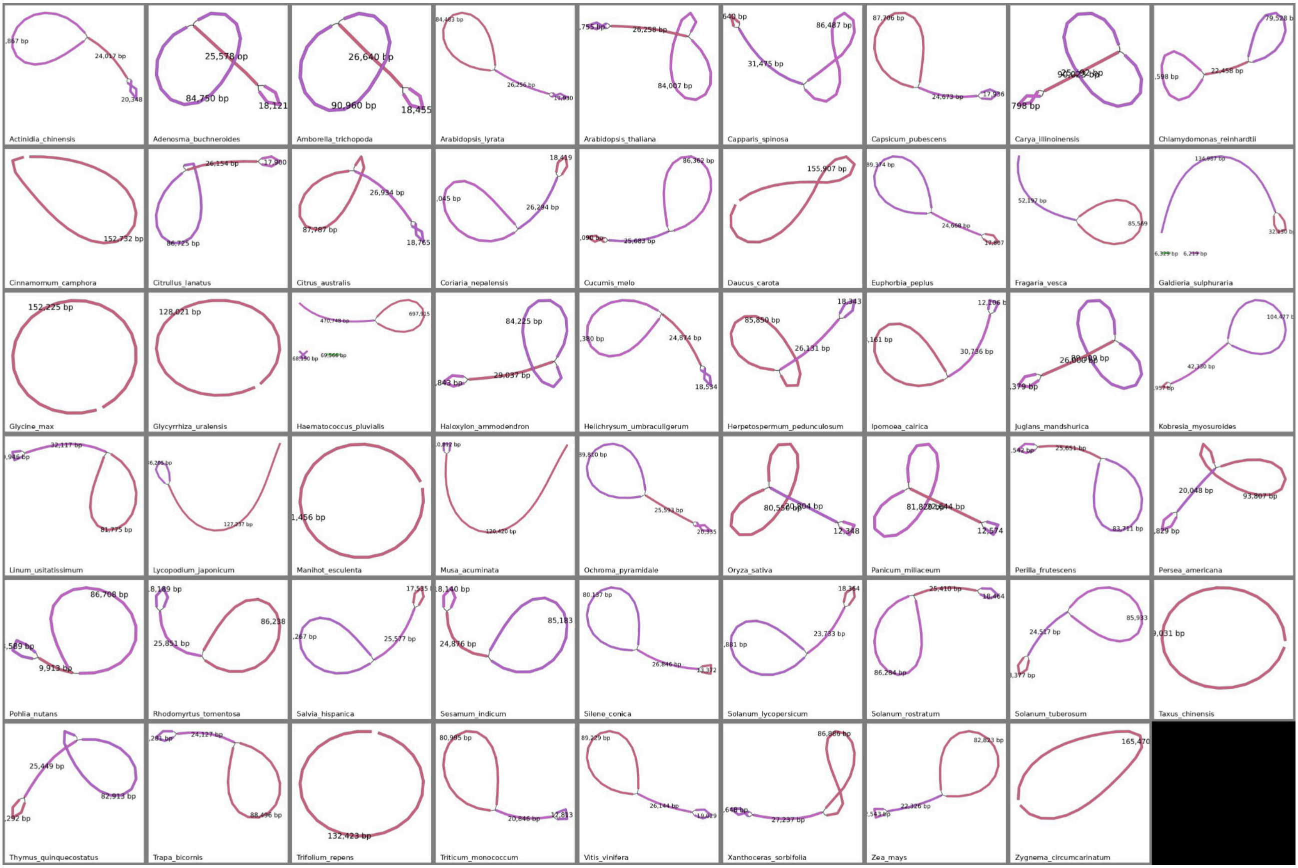


Figure S23: Chloroplast assembly graph of TIPPo. The coverage is set to 0.5x for each species, except for 0.25x for *T. chinensis,* 1.25x for *S. conica* and 0.075x for *L. japonicum.*
